# IL-36γ Drives Th9 cell Differentiation via IκBζ to Sustain Antitumor Immunity in Adoptive Cell Therapy

**DOI:** 10.64898/2026.04.29.721605

**Authors:** Qitai Zhu, Yu Chen, Jianling Gao, Peijun Tang, Zhenghao Ma, Xinying Li, Haolin Jiang, Ziyi Huang, Yachen Zang, Xin Zhao, Ji Zhang, Xuefeng Wang

## Abstract

Th9/Tc9 cells polarized with TGF-β and IL-4 exhibit superior antitumor efficacy, prompting extensive efforts to optimize their differentiation protocols and augment therapeutic potency. In the current study, we identified IL-36γ as a potent cytokine that synergizes with TGF-β to robustly drive Th9 differentiation both in the presence and absence of IL-4. IL-36γ-programmed Th9 cell subsets exhibited phenotypic and transcriptional profiles identical to classic Th9 cells. Mechanistically, IL-36γ drove Th9 cell differentiation through amplifying key signaling pathways such as STAT6, STAT5, and NF-κB that are essential for classic Th9 programming. Notably, we uncovered a novel regulatory axis wherein IL-36γ upregulates the transactivation factor IκBζ, which directly governs Th9 lineage specification. In in adoptive cell therapy (ACT) models, IL-36γ-polarized Th9 cell subsets demonstrated enhanced antitumor efficacy, attributable to their sustained persistence, reduced exhaustion markers and stem-like/memory properties. Collectively, this study elucidates a previously unrecognized IκBζ-dependent mechanism underpinning Th9 differentiation and highlights the translational potential of IL-36γ-engineered Th9 cells as a valuable ACT strategy for refractory tumors.

## Introduction

ACT that encompasses chimeric antigen receptor-T (CAR-T) cells, T cell receptor (TCR)-engineered T cells and tumor-infiltrating lymphocytes (TILs), has emerged as a transformative strategy for cancer immunotherapy^1–3^. While CD8^+^ cytotoxic T lymphocytes (CTLs) remain the primary focus, emerging evidence highlights the critical antitumor roles of CD4^+^ T helper cells^4^. Beyond their helper functions in orchestrating CTL responses, CD4^+^ T cells can directly eliminate MHC class II-expressing tumor cells and indirectly target MHC-deficient malignancies^5–7^.

Naïve CD4^+^ T cells differentiate into functionally distinct subsets including Th1, Th2, Th9, Th17 and induced regulatory T (iTreg) cell subsets under cytokine-specific polarization. Among these subsets, Th1 cells exhibit cytotoxic potential but suffer from rapid exhaustion in ACT^8,9^, whereas Th17 cells display prolonged persistence but incomplete effector maturation^10^. In contrast, Th9 cells uniquely combine sustained persistence, full cytolytic activity and resistance to exhaustion, positioning them as optimal candidates for ACT^10,11^. Classic Th9 cell polarization *in vitro* is characterized by co-expression of IL-9 alongside IL-10, IL-17, IL-21, and IL-22^12,13^. This process is orchestrated by two core signaling axes, TGF-β/PU.1 and IL-4/STAT6/IRF4^14–16^. While indispensable for Th9 generation, these pathways are not Th9-exclusive, as they concurrently regulate the differentiation of other Th subsets, highlighting the need to identify a master transcriptional regulator uniquely specifying Th9 identity. To date, studies reveal that Th9 polarization also relies on a collaborative network of transcription factors, including Id1, STAT5, ERG, DBP, NF-κB, BATF3, HIF1α, and Foxo1^14,17–26^, yet the definitive regulator governing Th9 lineage commitment remains elusive.

The antitumor efficacy of Th9 cells in ACT stems from the mechanisms as follows: IL-9-mediated enhancement of endogenous T cell and dendritic cell (DC) survival; IL-21-driven induction of IFN-γ production in endogenous CD8^+^ T and NK cells; and Granzyme B-dependent direct tumor cytotoxicity^27^. Critically, Th9 cells exhibit a functionally optimized profile characterized by reduced exhaustion markers, full cytotoxic capacity and enhanced proliferative potential, collectively enabling robust eradication of advanced tumors^10^.

Harnessing the therapeutic potential of Th9 cells in ACT requires efficient *in vitro* differentiation protocols. Conventional Th9-polarizing conditions achieve suboptimal induction efficiency^28,29^, prompting exploration of alternative stimuli. Notably, TNF-α potently enhances Th9 development *via* TNFR2-dependent activation of STAT5 and NF-κB^21^. OX40 costimulation drives Th9 differentiation through the TRAF6/NIK/NF-κBp52-RelB axis^29^. IL-1β amplifies IL-9/IL-21 secretion by Th9 cells *via* STAT1 phosphorylation and subsequent IRF1-mediated transcriptional activation of *Il9* and *Il21* promoters^28^. Intriguingly, IL-1β synergizes with IL-4 to induce IL-9 production comparable to classic TGF-β and IL-4 conditions, even in the absence of TGF-β^22^, underscoring its potency as a Th9-polarizing agent.

IL-36 agonists, including IL-36α, β and γ, are members of the IL-1 family and share the same receptor complex comprising IL-36 receptor (IL-36R) and IL-1RAcP^30^. Constitutively expressed on CD4^+^ T cells, IL-36R mediates Th1 cell differentiation *in vitro* and enhances Th1-driven immunity against Bacillus Calmette-Guerin (BCG) infection *in vivo*^31,32^. Intriguingly, IL-36γ upregulates IL-9 production across multiple CD4^+^ subsets including Th0, Th2, Th9 and induced regulatory T cells (iTregs) while simultaneously suppressing iTreg differentiation^33^. Under Th0 conditions, IL-36γ induces IL-9 production *via* NF-κBp50, IL-2/STAT5 and IL-4/STAT6 signaling pathways^33^, the core mechanisms conserved in classic Th9 cell programming^10,15,26^.

The pleiotropic nature of IL-36γ in modulating CD4^+^ T cell biology raises critical questions about whether IL-36γ-induced IL-9-producing CD4^+^ T cells under Th9 conditions exhibit distinct phenotypic signatures and antitumor efficacy compared to classic Th9 cells. Furthermore, the identity of the potential unique transcriptional driver specifically governing IL-36γ-mediated Th9 differentiation remains unresolved. To address these gaps, we aim to systematically characterize the molecular and functional profiles of IL-36γ-induced Th9 cells and delineate the novel mechanistic circuitry through which IL-36γ specifies Th9 lineage commitment.

## Results

### 1. IL-36γ synergizes with TGF-β to orchestrate Th9 differentiation both in the presence and absence of IL-4 while preserving classic Th9 transcriptional programs

Classic Th9 cell induction relies on TGF-β and IL-4. Given that IL-36γ enhances IL-9 expression^33^, we sought to investigate whether IL-36γ can induce IL-9-overexpressing Th9 cells independently of TGF-β, IL-4 or both of them. OT-II naïve CD4^+^ T cells were primed with OVA_323-339_ peptide-loaded antigen-presenting cells (APCs) for 2 days in the presence of TGF-β, IL-4, IL-36γ or in combination. Flow cytometry and enzyme-linked immunosorbent assay (ELISA) revealed minimal IL-9 production under single-cytokine treatments (Figure 1A and 1B). As expected, TGF-β plus IL-4 induced robust expression of IL-9, which was further amplified by IL-36γ supplementation (Figure 1A and 1B), consistent with prior findings^33^. Strikingly, IL-36γ in combination with TGF-β elicited substantial levels of IL-9 even in the absence of IL-4, which was comparable to that produced by classic Th9 cells (Figure 1A and 1B). Meanwhile, IL-10 secretion was also significantly increased by IL-36γ in combination with TGF-β both in the presence and absence of IL-4 (Figure S1A). Additionally, dose-response assays confirmed that IL-36γ potentiated IL-9 production under classic Th9 polarization in a concentration-dependent manner (Figure 1C and S1B). These results establish IL-36γ as a potent agent that synergizes with TGF-β to drive IL-9 production in primed CD4^+^ T cells both in the presence and absence of IL-4.

**Figure 1.**
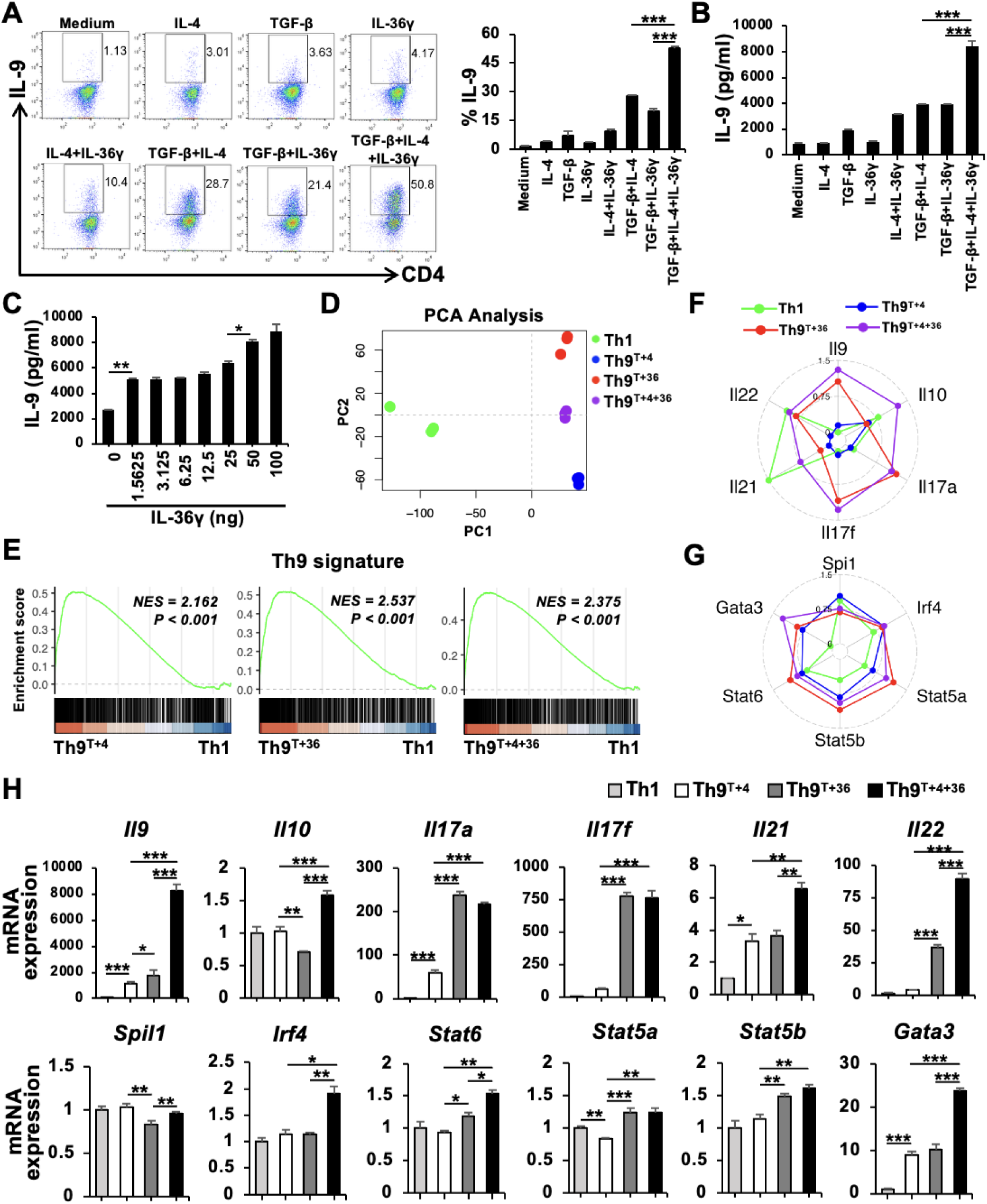
IL-36γ drives Th9 differentiation while preserving classic Th9 transcriptional programs. (A and. **B)** Naïve CD4^+^ T cells from OT-II mice were activated with OVA_323-339_ peptide-pulsed APCs and cultured in the presence of TGF-β (2.5 ng/ml), IL-4 (10 ng/ml), IL-36γ (100 ng/ml) or in indicated combinations for 2 days. (**A)** Intracellular IL-9 expression in CD4^+^ T cells was quantified by flow cytometry (left, representative plots; right, quantification). (**B)** IL-9 levels in culture supernatants were measured *via* ELISA. **(C)** Naïve CD4^+^ T cells from OT-II mice were activated with OVA_323-339_-pulsed APCs and cultured under classic Th9-polarizing conditions supplemented with graded concentrations of IL-36γ for 2 days. IL-9 production in culture supernatants was quantified by ELISA. **(D-H)** Transcriptomic profiling of CD4^+^ T cells that were activated by plate-bound anti-CD3/CD28 mAbs (each 5 μg/ml) in the presence of IL-2 (20 ng/ml) and polarized for 36h under Th1, Th9^TGF-β+IL-4^ (Th9^T+4^), Th9^TGF-β+IL-36γ^ (Th9^T+4^) or Th9^TGF-β+IL-4+IL-36γ^ (Th9^T+4+36^) conditions. **(D)** PCA of global transcriptomes across T cell subsets. **(E)** GSEA against Th9-defining signatures (GSE222909), demonstrating conserved lineage identity in IL-36γ-prgrammed Th9 subsets compared to Th1 controls. **(F and G)** Radar plots illustrating expression patterns of classic Th9-associated cytokines **(F)** and classic Th9 lineage-determining transcription factors **(G)** across T cell subsets as indicated. **(H)** qRT-PCR validation of key transcriptional markers showing high concordance with RNA-SEQ profiles. For all *in vitro* experiments, one-way ANOVA was used **(A-C and H)**. All statistical tests were two-sided and all replicates were biologically independent samples (*n* = 3). \**p* < 0.05, \*\**p* < 0.01, \*\*\**p* < 0.001.

Given the broad expression of IL-36R across T cells, monocytes and dendritic cells (DCs), we next assessed whether IL-36γ directly regulates IL-9 production in CD4^+^ T cells. Purified naïve CD4^+^ T cells were activated with anti-CD3/CD28 mAbs in the presence of TGF-β, IL-4, IL-36γ or in combination for 2 days. Flow cytometric analysis revealed that consistent with prior observations, IL-36γ significantly amplified IL-9 production under classic Th9-polarizing conditions (Figure S1C and S1D). Strikingly, in the absence of IL-4, IL-36γ combined with TGF-β induced IL-9 production surpassing those of classic Th9 cells and approaching the output achieved by IL-36γ-supplemented classic Th9 conditions (Figure S1C and S1D). These findings demonstrate that IL-36γ directly programs IL-9 expression in CD4^+^ T cells through a TGF-β-dependent yet IL-4-independent mechanism.

In view of the evidence that IL-36 signaling amplifies Th1 responses^31,34^, we investigated whether IL-36γ-induced Th9 cells retained transcriptional hallmarks of classic Th9 cells rather than adopting Th1-like features. RNA sequencing (RNA-SEQ) was performed on 4 subsets: TGF-β plus IL-36γ-induced Th9 (Th9^TGF-β+IL-36γ^) cells, IL-36γ-elicited Th9 (Th9^TGF-β+IL-4+IL-36γ^) cells under classic Th9 conditions, classic Th9 (Th9^TGF-β+IL-4^) cells and Th1 cells. Principal component analysis (PCA) revealed that both Th9^TGF-β+IL-36γ^ and Th9^TGF-β+IL-4+IL-36γ^ subsets exhibited transcriptional proximity to classic Th9 cells, with clear divergence from Th1 cells (Figure 1D). A three-way comparison of overlapping transcripts identified 801 shared genes between Th9^TGF-β+IL-4+IL-36γ^ cells and classic Th9^TGF-β+IL-4^ cells, 779 shared genes between Th9^TGF-β+IL-36γ^ cells and classic Th9^TGF-β+IL-4^ cells and 638 core genes conserved across all three Th9 subsets (Figure S1E).

To evaluate whether Th9^TGF-β+IL-36γ^ cells and Th9^TGF-β+IL-4+IL-36γ^ cells retain classic Th9 characteristics, we performed gene set enrichment analysis (GSEA) against a predefined classic Th9 transcriptional signature (GSE222909)^18^. The analysis revealed significant enrichment of Th9-defining genes in both IL-36γ-polarized subsets compared to Th1 cells, confirming their transcriptional fidelity to classic Th9 lineages (Figure S1E).

To rigorously assess whether Th9^TGF-β+IL-36γ^ and Th9 ^TGF-β+IL-4+IL-36γ^ cells harbor classic Th9-defining features, we analyzed their transcriptional profiles in triplicate based on RNA-SEQ data. Strikingly, these IL-36γ-polarized Th9 cells exhibited robust maintenance or upregulation of core cytokines (*Il9, Il10, Il17a, Il21 and Il22*) and signature transcription factors (*Spi1, Irf4, Stat5, Stat6 and Gata3*), all hallmarks of classic Th9 identity (Figure 1F and 1G). The quantitative reverse transcription-PCR (qRT-PCR) validation confirmed high concordance with RNA-SEQ findings (Figure 1H), solidifying the transcriptional consistency of IL-36γ-induced Th9 subsets to their classic counterparts.

Collectively, these data establish that both in the presence and absence of IL-4, IL-36γ synergizes with TGF-β to robustly induce Th9 cells with phenotypic and transcriptional profiles identical to classic Th9 cells.

### 2. IL-36γ drives Th9 cell differentiation through enhancing STAT6, STAT5 and NF-κB signaling, core pathways governing classic Th9 programming

To delineate the molecular circuitry underlying IL-36γ-driven Th9 differentiation, we systematically profiled key transcriptional regulators governing Th9 lineage commitment. Integrated analysis of RNA-SEQ and qRT-PCR datasets revealed upregulation of *Stat6*, *Il2*, *Stat5a*, *Stat5b* and *Nfκb2* in both IL-36γ-polarized Th9 subsets (Figure 2A, 2B and 1H), implicating these pathways mediating IL-36γ-driven Th9 cell differentiation.

**Figure 2.**
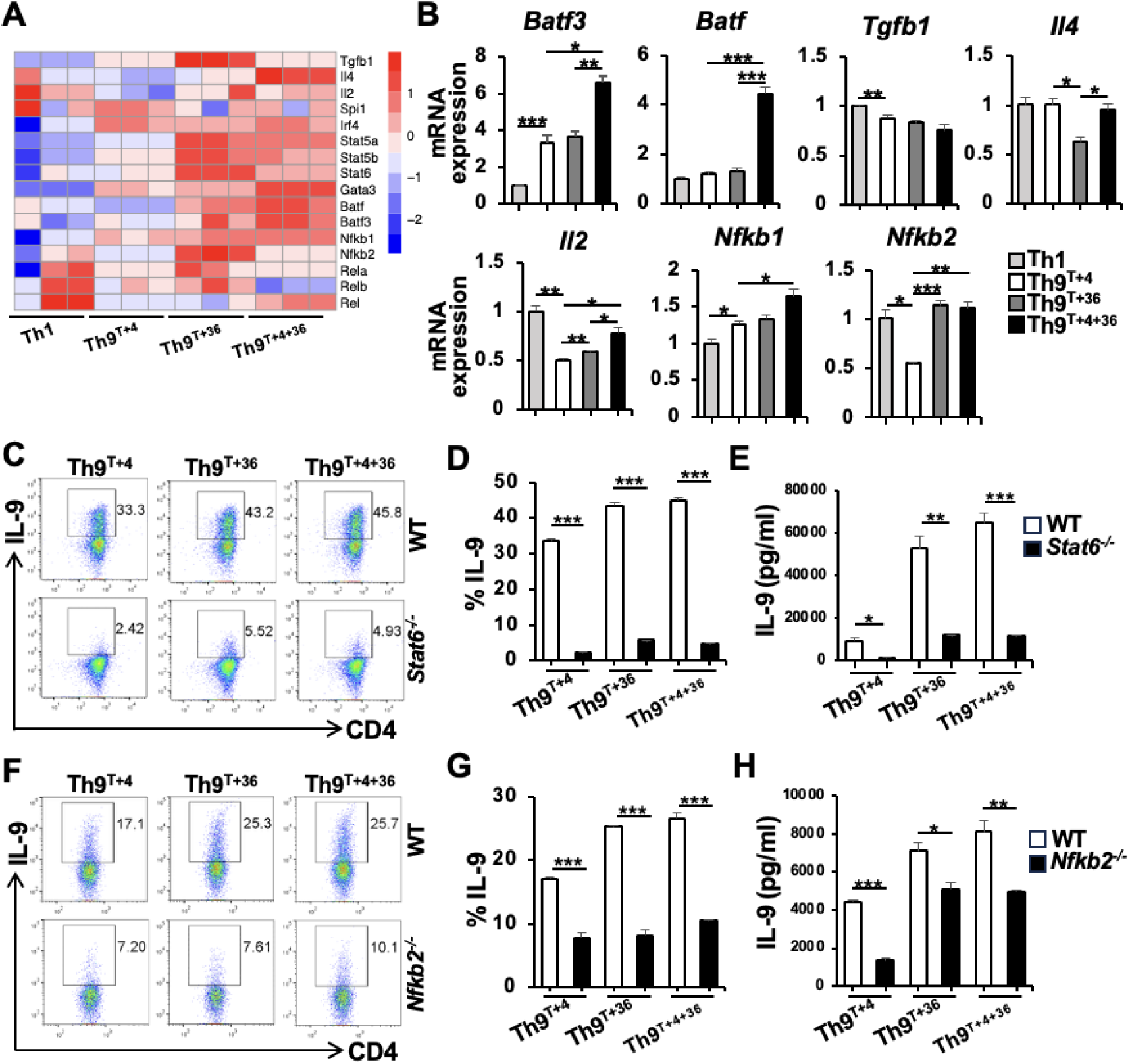
IL-36γ drives Th9 cell differentiation through enhancing STAT6 and NF-κB2 signaling pathways. **(A)** Heatmap based on RNA-SEQ data showing expression profiles of classic Th9-associated genes in *in vitro*-polarized Th1, Th9^TGF-β+IL-4^, Th9^TGF-β+IL-36γ^ and Th9^TGF-β+IL-4+IL-36γ^ subsets. **(B)** qRT-PCR validation of transcriptional levels of indicated cytokines and transcription factors across Th1, Th9^TGF-β+IL-4^, Th9^TGF-β+IL-36γ^ and Th9^TGF-β+IL-4+IL-36γ^ subsets. **(C-E)** Naïve CD4^+^ T cells from WT and *Stat6*^-/-^ mice were respectively activated with plate-bound anti-CD3/CD28 mAbs (each 5 μg/ml) and IL-2 (20 ng/ml) and cultured under Th9^TGF-β+IL-4^, Th9^TGF-β+IL-36γ^ or Th9^TGF-β+IL-4+IL-36γ^ conditions for 2 days. **(C)** Representative flow cytometry plots of intracellular IL-9^+^ cells. **(D)** Quantitative analysis of IL-9^+^ cell frequencies. **(E)** IL-9 concentrations in culture supernatants measured by ELISA. **(F-H)** Naïve CD4^+^ T cells from WT and *Nfkb2*^-/-^ mice were respectively activated with anti-CD3/CD28 mAbs (each 5 μg/ml) and IL-2 (20 ng/ml) and cultured under Th9^TGF-β+IL-4^, Th9^TGF-β+IL-36γ^ or Th9^TGF-β+IL-4+IL-36γ^ conditions for 2 days. **(F)** Representative flow cytometry plots of intracellular IL-9^+^ cells. **(G)** Quantitative analysis of IL-9^+^ cell frequencies. **(H)** IL-9 concentrations in culture supernatants measured by ELISA. One-way ANOVA was used for comparisons of four groups**(B)**. Comparisons between WT and Stat6^-/-^ or *Nfkb2*^-/-^groups within the same Th9 subset were performed using unpaired Student’s *t*-test **(D, E, G and H)**. All statistical tests were two-sided and all replicates were biologically independent samples (*n* = 3). \**p* < 0.05, \*\**p* < 0.01, \*\*\**p* < 0.001.

To validate the functional relevance of these regulators to IL-36γ-mediated Th9 induction, we employed chemical inhibitors and/or genetically deficient CD4^+^ T cells targeting key pathway components. First, wild-type (WT) and *Stat6^-/-^* CD4^+^ T cells were polarized under classic Th9 conditions or IL-36γ-containing regimens. As expected, STAT6 deficiency abolished classic Th9 differentiation (Figure 2C-2E), consistent with its established role in IL-9 regulation^15^. Interestingly, both Th9^TGF-β+IL-36γ^ and Th9^TGF-β+IL-4+IL-36γ^ subsets were almost abrogated in *Stat6*^-/-^ CD4^+^ T cells (Figure 2C-2E), establishing STAT6 as a non-redundant element for IL-36γ-driven Th9 polarization^33^. Pharmacological inhibition using AS1517499 (selective STAT6 inhibitor) dose-dependently suppressed IL-36γ-induced Th9 generation (Figure S2A and S2B), further corroborating this mechanism^33^.

BATF and BATF3, downstream effectors of the IL-4/STAT6 signaling^35^, are known regulators of IL-9 secretion in classic Th9 polarization^23^. RNA-SEQ and qRT-PCR analyses revealed that *Batf* and *Batf3* were upregulated in IL-36γ-induced Th9 cells (Figure 2A and 2B), prompting further investigation into their functional roles. shRNA-mediated knockdown of *Batf3* but not *Batf* (data not showed) selectively attenuated IL-9 production in Th9^TGF-β+IL-4+IL-36γ^ cells (Figure S2C and S2D), indicating a non-redundant role for BATF3 in this context.

In that IL-36γ-mediated enhancement of the IL-2/STAT5 signaling pathway (Figure 2A, 2B and 1H), we next assessed its functional necessity in Th9 differentiation. Pretreatment with 573108-M, a selective STAT5 inhibitor, elicited dose-dependent suppression of IL-9 production in both Th9^TGF-β+IL-36γ^ and Th9^TGF-β+IL-4+IL-36γ^ subsets, paralleling its inhibitory effects on classic Th9 cells (Figure S2E and S2F). These findings definitively establish the IL-2/STAT5 signaling as an essential mediator of IL-36γ-driven Th9 differentiation.

The NF-κB pathway is a well-established prerequisite for Th9 cell differentiation^10,21,22,36,37^. Interestingly, in addition to the fact that IL-36 agonists effectively active NF-κB pathway^38^, our data demonstrated that IL-36γ also increased the transcript levels of *Nfκb1* in Th9^TGF-β+IL-4+IL-36γ^ cells and *Nfκb2* levels in both Th9^TGF-β+IL-36γ^ and Th9^TGF-β+IL-4+IL-36γ^ cells, compared to classic Th9 cells (Figure 2A and 2B).

Functional validation using QNZ, a potent NF-κB inhibitor, demonstrated dose-dependent suppression of IL-9 production across IL-36γ-polarized and classic Th9 subsets (Figure S2G and S2H), conclusively linking IL-36γ-indcued Th9 cell differentiation to amplification of NF-κB signaling.

While the non-canonical RelB/NF-κBp52 pathway but not the canonical RelA/NF-κBp50 axis has been shown to directly drive *Il9* transcription in OX40-stimulated Th9 cells^29^, our findings reveal a broader role for this pathway. Specifically, IL-36γ significantly upregulated *Nfκb2* (encoding the p100 precursor of NF-κBp52) in both Th9^TGF-β+IL-36γ^ and Th9^TGF-β+IL-4+IL-36γ^ subsets (Figure 2A and 2B), prompting investigation into its functional necessity. Strikingly, NF-κBp52 deficiency severely impaired IL-9 production in IL-36γ-polarized Th9 cells (Figure 2F-2H) and partially disrupted classic Th9 differentiation, a phenotype not previously resolved^10,29^. These results unequivocally demonstrate that IL-36γ amplifies the RelB/NF-κBp52 signaling pathway to license Th9 cell expansion. Intriguingly, while intracellular IL-9 staining by flow cytometry showed a 3.1-fold reduction in *Nfkb2*^-/-^ Th9^TGF-β+IL-36γ^ cells and 2.5-fold reduction in *Nfkb2*^-/-^ Th9^TGF-β+IL-4+IL-36γ^ cells (Figure 2F and 2G), secreted IL-9 levels quantified by ELISA demonstrated only 1.4-fold decrease in *Nfkb2*^-/-^ Th9^TGF-β+IL-36γ^ cells and 1.6-fold decrease in *Nfkb2*^-/-^ Th9^TGF-β+IL-4+IL-36γ^ cells (Figure 2H), suggesting that NF-κBp52 predominantly exerts an effect on differentiation of IL-36γ-polarized Th9 cells during the late-phase.

Together, these data conclusively demonstrate that STAT6, STAT5 and NF-κB pathways, previously established as core regulators of classic Th9 cell generation^10,15,18^, synergistically orchestrate IL-36γ-driven Th9 differentiation through conserved yet amplified signaling nodes.

### 3. IκBζ emerged as a critical transactivation factor that orchestrated IL-36γ-driven Th9 cell differentiation

While IL-36γ-induced Th9 cell differentiation was characterized through conventional pathways, the identification of a unique transcriptional regulator driving this process remains elusive. The NF-κB pathway serves as the primary downstream effector of IL-36/IL-36R signaling^38^. Upon IL-36 stimulation, cytoplasmic IκBs undergo phosphorylation and degradation, enabling NF-κB nuclear translocation. In contrast, nuclear atypical IκBs (Bcl-3, IκBζ, IκBNS, IκBη and IκBL) modulate NF-κB activity by either suppressing transcriptional programs or enhancing specific gene expression^39^. Interestingly, RNA-SEQ analysis identified that *Nfκbiz* (encoding IκBζ) rather than other atypical IκBs was upregulated in both IL-36γ-polarized Th9 subsets versus classic Th9 cells (Figure 3A and 3B). Subsequent qRT-PCR confirmed elevated *Nfκbiz* mRNA in both IL-36γ-induced Th9 subsets (Figure 3B), with parallel increase in IκBζ protein at different time points post-differentiation (Figure 3C, S3A and S3B).

**Figure 3.**
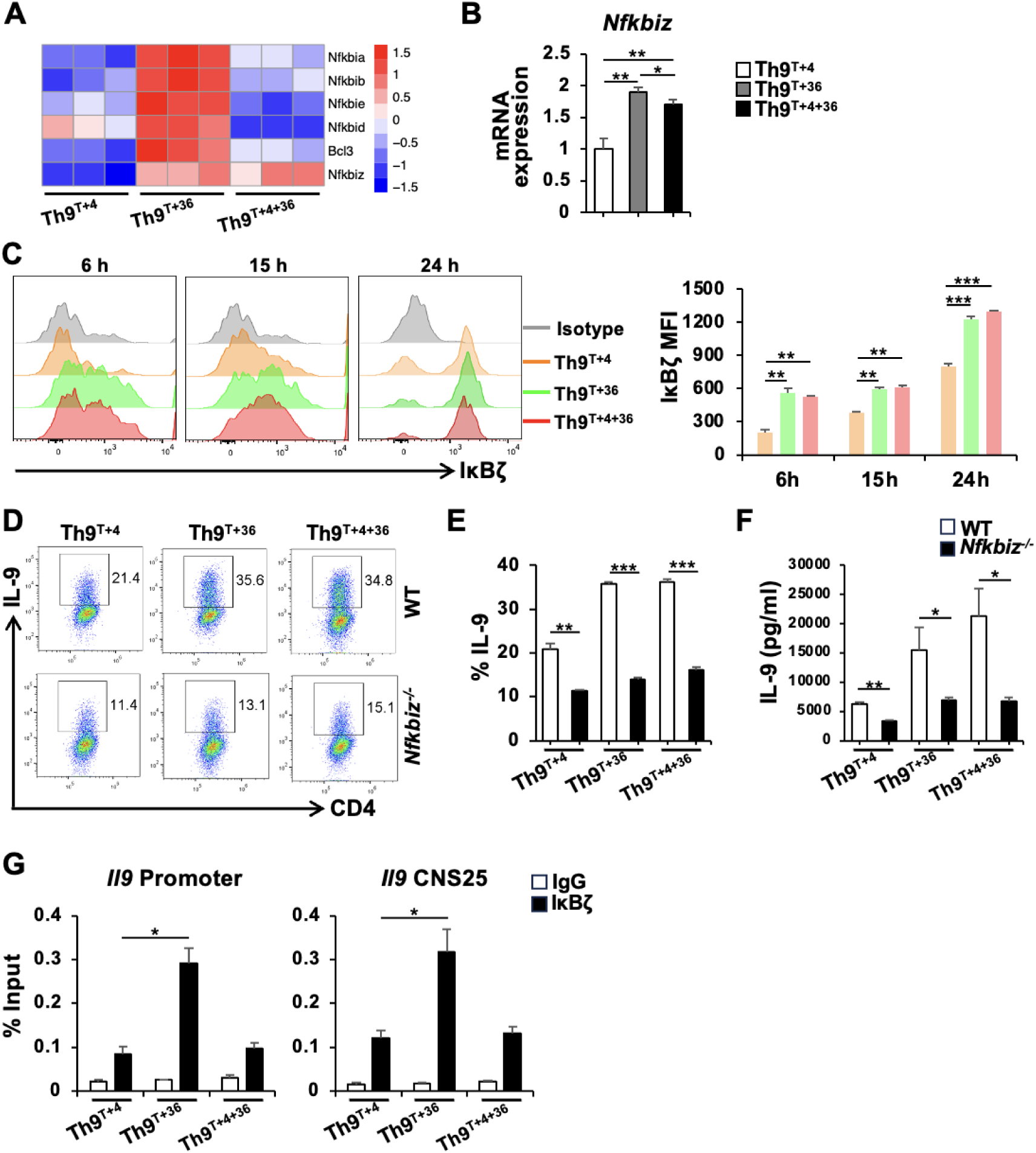
IκBζ licenses IL-36γ-enhanced Th9 differentiation by orchestrating IL-9 transcriptional activation. **(A)** Heatmap based on RNA-SEQ data depicting transcriptional profiles of IκBs in Th9^TGF-β+IL-4^, Th9^TGF-β+IL-36γ^ or Th9^TGF-β+IL-4+IL-36γ^ subsets. **(B)** *Nfkbiz* mRNA levels quantified by qRT-PCR across indicated Th9 subsets. **(C)** Naïve CD4^+^ T cells were activated with anti-CD3/CD28 mAbs (each 5 μg/ml) and IL-2 (each 20 ng/ml) and cultured under different Th9 subset-polarizing conditions as indicated. Cells were harvested at 6, 15 and 24 h. Time-dependent IκBζ expression was quantified by flow cytometry (left, representative histograms; right, quantification). **(D-F)** Naïve CD4^+^ T cells were polarized under different Th9-inducing conditions as **c** for 2 days. **(D)** Representative flow cytometry plots of IL-9^+^ cells. **(E)** Frequency of IL-9^+^ T cells. **(F)** IL-9 levels in supernatants measured by ELISA. **(G)** Naïve CD4^+^ T cells were polarized under different Th9-inducing conditions as **c** for 36 h. ChIP-qPCR analysis of IκBζ binding to *Il9* promoter and CNS25 enhancer regions across indicatedTh9 subsets. The data **(B, C and G)** were statistically analyzed using one-way ANOVA. An unpaired Student’s *t*-test was employed for comparisons between WT and IκBζ knockout groups within the same Th9 subset **(E and F)**. All statistical tests were two-sided and all replicates were biologically independent samples (*n* = 3). \**p* < 0.05, \*\**p* < 0.01, \*\*\**p* < 0.001.

Next, the functional role of IκBζ was assessed using *Nfκbiz*^-/-^ CD4^+^ T cells. Intriguingly, IκBζ deficiency reduced classic Th9 differentiation by >50% (Figure 3D-3F). Strikingly, IL-36γ-programmed Th9 subsets exhibited even greater differentiation impairment (Figure 3D-3F). Critically, the genetic ablation of *Nfκbiz* abolished the IL-36γ-driven Th9 expansion, rendering IL-36γ-induced Th9 frequencies almost indistinguishable from that of classic Th9 cells (Figure 3D-3F), implicating IκBζ as a central mediator in IL-36γ promoting Th9 differentiation.

To investigate whether IκBζ directly promotes Th9 cell differentiation, we performed chromatin immunoprecipitation (ChIP) assays to assess its binding to the *Il9* promoter and enhancer^40^. Our data revealed that IκBζ robustly bound to the IL-9 promoter and CNS-25 enhancer across all Th9 differentiation models. Meanwhile, Th9^TGF-β+IL-36γ^ but not Th9^TGF-β+IL-4+IL-36γ^ cells displayed significantly stronger IκBζ enrichment at these loci versus classic Th9 cells (Figure 3G), establishing direct transcriptional regulation of Il9 by IκBζ.

Therefore, IκBζ as the master regulator of IL-36γ-driven Th9 cell differentiation, demonstrating its necessity for enhanced IL-9 production via direct chromatin binding to the *Il9* locus, and revealing its non-redundant role in amplifying Th9 differentiation beyond canonical pathways.

### 4. IL-36γ-induced Th9 cell subsets exhibited superior antitumor efficacy with prolonged engraftment in adoptive therapy

To evaluate the therapeutic efficacy of IL-36γ-induced Th9 cell subsets, we developed an ACT model. B16-OVA melanoma-bearing mice (CD45.2^+^) received CTX or PBS pretreatment on day 5 post-tumor inoculation, followed by adoptive transfer of OVA-specific CD45.1^+^Th1, CD45.1^+^Th9^TGF-β+IL-4^, CD45.1^+^Th9^TGF-β+IL-36γ^ or CD45.1^+^Th9^TGF-β+IL-4+IL-36γ^ subsets, respectively combined with OVA_323-339_-pulsed bone marrow-derived dendritic cells (BMDCs) on day 6 (Figure 4A). Intriguingly, all Th9 subsets demonstrated superior therapeutic efficacy over Th1 cells (Figure 4A). Notably, Th9^TGF-β+IL-4+IL-36γ^ cells exhibited remarkably enhanced antitumor activity compared to classic Th9 cells, while Th9^TGF-β+IL-36γ^ showed comparable efficacy to classic Th9 cells (Figure 4A). Survival analysis further demonstrated that mice receiving both Th9^TGF-β+IL-4+IL-36γ^ and Th9^TGF-β+IL-36γ^ cell therapies achieved significant extension compared to classic Th9 cells (Figure 4B). These findings collectively demonstrate that IL-36γ-induced Th9 cells exhibit more potent tumor inhibition compared to classic Th9 cells in ACT.

**Figure 4.**
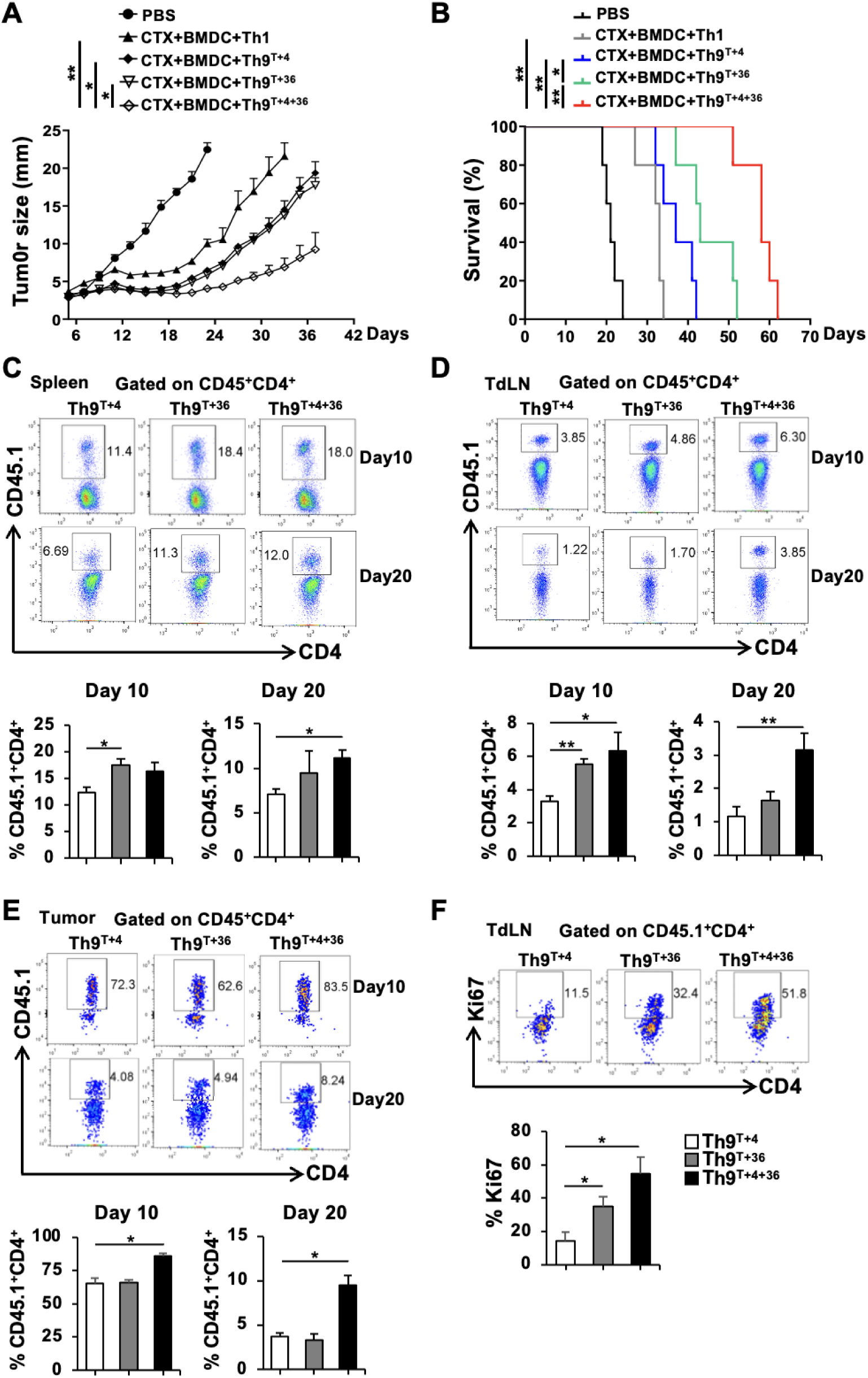
IL-36γ-programmed Th9 cells exhibit enhanced tumor control and sustained persistence in adoptive cell therapy. **(A)** Comparative analysis of tumor growth kinetics in B16-OVA-bearing mice (*n* = 5) subjected to different treatments as indicated. **(B)** Survival curves of tumor-bearing mice (*n* = 5) post-treatment. **(C-F)** Tumor-bearing mice were sacrificed on day 10 and 20 post-transfer for immunological profiling. Flow cytometric analysis of representative plots (up) and quantification (below) for adoptively transferred CD45.1^+^CD4^+^ T cells in spleens **(C)**, TdLNs **(D)** and tumors **(E). (F)** Flow cytometric analysis of representative plots (up) and quantification (below) for Ki67^+^ proliferation rates in TdLN-resident adoptively transferred CD45.1^+^CD4^+^ T cells on day 10. One-way ANOVA was employed for data analysis **(A and C-F)**. Kaplan-Meier survival curves were compared by Log-rank test **(B)**. \**p* < 0.05, \*\**p* < 0.01.

During tumor adoptive immunotherapy, the long-term persistence of transferred T cells is crucial for sustaining therapeutic efficacy. Subsequently, we analyzed the spatiotemporal distribution of adoptively transferred T cells as well as endogenous T cell subsets in tumor-bearing mice. First, CD45^+^ cell frequencies were found to be comparable in tumor tissues across three Th9 subset-treated cohorts on day 10 and 20 after adoptive transfer (Figure S4B). Meanwhile, IL-36γ-induced Th9 cell-treated groups had comparable CD4^+^ T cell frequencies in spleens, tumor-draining lymph nodes (TdLNs) and tumor tissues relative to classic Th9 cell regimen on days 10 and 20 post-transfer, except that a transient downregulation was observed in the Th9^TGF-β+IL-36γ^ cell-treated group on day 10, which resolved by day 20 (Figure S4C). Given the pivotal role of CTLs in directly eradicating tumors, we observed an increase in the frequency of CD8^+^ T cells on day 10 that diminished by day 20 (Figure S4C and S4D), suggesting that IL-36γ-programmed Th9 cells may enhance early-phase CD8^+^ T cell infiltration or proliferation in tumor and lymphoid organs.

Then, detailed analysis demonstrated that adoptively transferred Th9^TGF-β+IL-4+IL-36γ^ cells (CD45.1^+^CD4^+^) exhibited higher frequency in spleens, TdLNs and tumor tissues compared to classic Th9 cells (CD45.1^+^CD4^+^) on both day 10 and 20 (Figure 4C-4E). Interestingly, while Th9^TGF-β+IL-36γ^ cells showed comparable tumor infiltration to classic Th9 cells, it displayed elevated frequency in spleens and TdLNs on day 10 (Figure 4C-4E). These findings collectively indicate that IL-36γ-induced Th9 cells possess superior in *vivo* persistence compared to classic Th9 cells, with this advantage persisting into the late therapeutic phase.

In view of superior persistence of IL-36γ-induced Th9 cells, we evaluated Ki67 expression profiles of transferred T cell subsets on day 10 after transfer. Interestingly, both IL-36γ-induced Th9 cell subsets significantly upregulated Ki67 compared to classic Th9 cells within TdLNs (Figure 4F). This proliferative advantage was also pronounced in Th9^TGF-β+IL-36γ^ subset in spleens compared to classic Th9 cells (Figure S4E). Intriguingly, despite no differences in Ki67 expression across three kinds of tumor-infiltrating Th9 cell subsets, all exhibited high levels of Ki67 (Figure S4F), suggesting that they harbored robust proliferative activity within the tumor niche. Additionally, Gene Ontology (GO) enrichment analysis revealed that genes upregulated in IL-36γ-programmed Th9 subsets compared to classic Th9 cells were predominantly associated with T cell proliferation, activation and lineage commitment (Figure S4G).

These findings collectively demonstrate that IL-36γ-induced Th9 cells acquire enhanced clonal expansion capacity post-transfer, enabling prolonged persistence and spatial dominance that mechanistically underpin their superior antitumor efficacy.

### 5. IL-36γ-induced Th9 cells are distinct effectors with full cytotoxicity

Because IL-36γ-programmed Th9 cells are promising T cell lineage for ACT, we sought to characterize their antitumor effector profiles. First, GSEA data revealed that both Th9^TGF-β+IL-4+IL-36γ^ and Th9^TGF-β+IL-36γ^ subsets significantly upregulated Th1-associated gene signatures (GSE22886) compared to classic Th9 cells (Figure 5A). T Cell State Identifier (TCellSI) is a novel tool that systematically evaluates T cell signatures by calculating the T Cell State Score^41^. Then, the cytotoxicity scoring calculated by TCellSI tool demonstrated elevated cytotoxic potential in IL-36γ-modified Th9 subsets (Figure 5B), corroborating the enhanced effector T cell gene features (GSE41867) (Figure 5C). Furthermore, heatmap analysis of RNA-seq data identified that IL-36γ-programmed Th9 subsets had distinct effector cytokine profiles. It showed upregulation of a series of effector cytokines (*Ifng*, *Gzma*, *Gzmc*, *Gzmk*, *Il6*, *Il24*, *Il17a* and *Il17f)* with *Gzmb* downregulation. *Tnf* exhibited context-dependent regulation, upregulated in Th9^TGF-β+IL-36γ^ cells but downregulated in Th9^TGF-β+IL-4+IL-36γ^ cells (Figure 5D). Costimulatory molecule profiling revealed the increased expression of *Tnfsf8* but downregulation of *Tnfsf14*, *Tnfrsf13b* and *Icos* in IL-36γ-induced two Th9 subsets, and upregulation of *Tnfrsf4*, *Tnfrsf9*, *Tnfrsf14* and *Tnfrsf18* in Th9^TGF-β+IL-36γ^ cells (Figure 5D). Next, qRT-PCR validation confirmed upregulation of *Ifng*, Gzma, Gzmc and Gzmk and downregulation of *Gzmb* in IL-36γ-modified two Th9 subsets, while *Tnf* remained unchanged (Figure 5E). In addition, transcriptional factor analysis demonstrated elevated *Stat4*, *Runx3* and *Tbx21* with reduced *Eomes* and *Stat1* in IL-36γ-polarized Th9 subsets (Figure S5A and 5B), defining a unique effector phenotype distinct from that of classic Th9 cells. These findings demonstrate that IL-36γ-polarized Th9 subsets exhibit pre-existing functional superiority over classic Th9 cells, manifesting enhanced antitumor effector profiles even prior to adoptive transfer into tumor-bearing mice.

**Figure 5.**
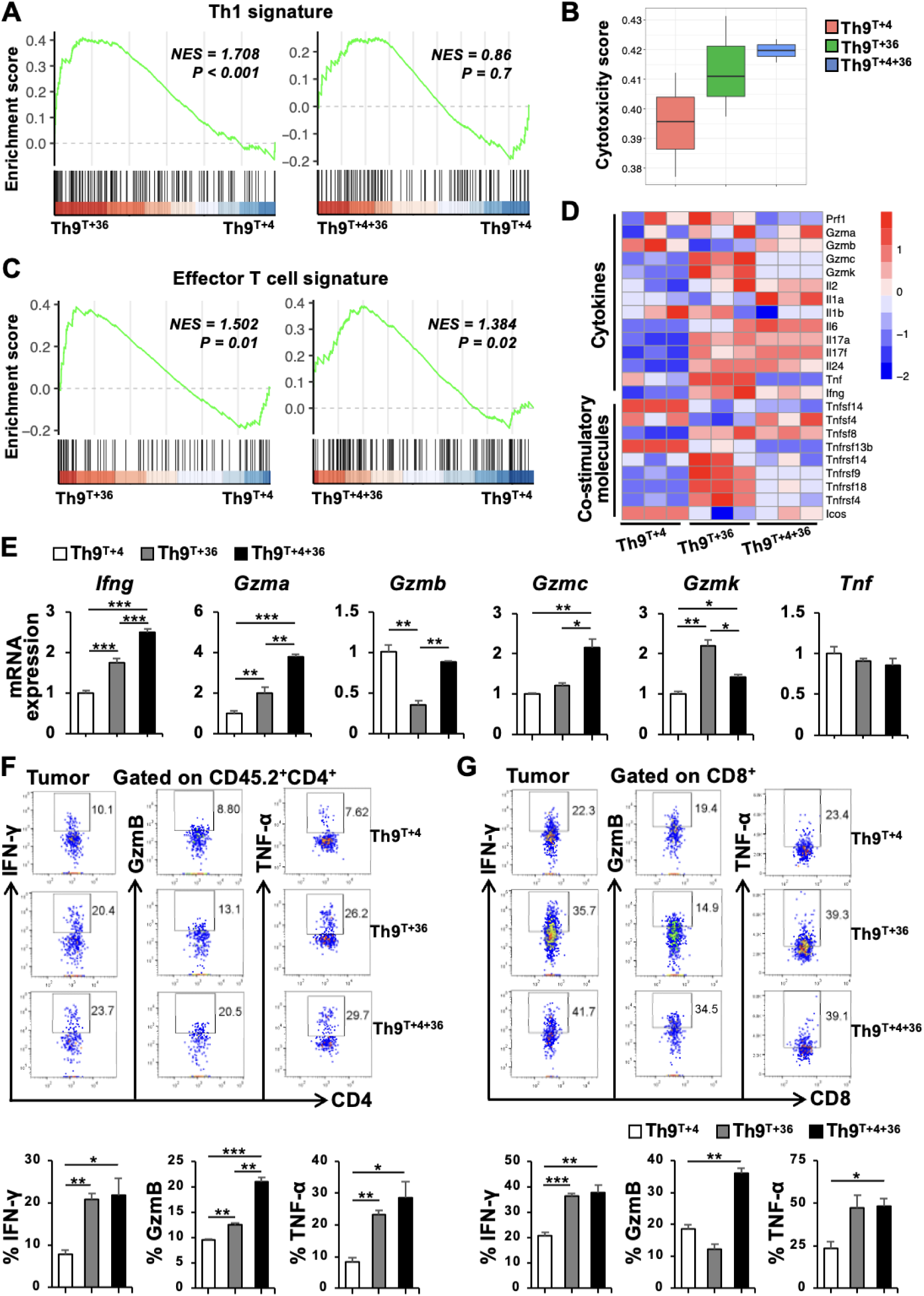
IL-36γ-programmed Th9 subsets are endowed with enhanced cytotoxic effector phenotypes and potentiate endogenous T cell antitumor responses post-transfer. **(A)** GSEA comparing Th9^TGF-β+IL-36γ^ and Th9^TGF-β+IL-4+IL-36γ^ subsets versus Th9^TGF-β+IL-4^ cells, showing enriched Th1 signature (GSE22886). **(B)** Cytotoxicity scores computed by TCellSI tool for Th9 subsets. **(C)** Effector T cell signature enrichment (GSE41867) across three Th9 subsets. **(D)** Heatmap based on RNA-SEQ data demonstrating differentially expressed cytotoxic and costimulatory molecules. **(E)** qRT-PCR validation of effector cytokine transcripts in three Th9 cells. **(F and G)** Flow cytometric profiling of IFN-γ^+^, GzmB^+^, and TNF-α^+^ endogenous CD45.2^+^CD4^+^ T **(F)** and CD45.2^+^CD8^+^ T **(G)** cells in tumors 10 days post-ACT (up, representative plots; below, quantification). One-way ANOVA was employed for data analysis **(E-G)**. \**p* < 0.05, \*\**p* < 0.01, \*\*\**p* < 0.001.

Given the effector-persistence dichotomy in functionally potent T cells and the immunomodulatory capacity of adoptively transferred CD4^+^ T cells on host tumor-infiltrating lymphocytes (TILs), we subsequently assessed the expression profiles of effector molecules in exogenous CD4^+^ T cells (CD45.1^+^), host-derived CD4^+^ T cells (CD45.2^+^) and CD8^+^ T cells after adoptive therapy. Unexpectedly, adoptively transferred CD4^+^ T cells across all cohorts receiving T cell therapy showed comparable levels of IFN-γ, GzmB and TNF-α, with the exception of elevated IFN-γ in the Th9^TGF-β+IL-36γ^ treatment cohort (Figure S5C), suggesting that IL-36γ-programmed Th9 subsets possess fully effector function as classic Th9 cells^10^. Strikingly, the tumors from mice receiving Th9^TGF-β+IL-4+IL-36γ^ and Th9^TGF-β+IL-36γ^ subsets exhibited pronounced upregulation of these effector molecules in endogenous CD4^+^ and CD8^+^ T cell populations (Figure 5F and 5G). Our data reveal a key mechanism that IL-36γ-polarized Th9 cells, though lacking intrinsic cytotoxic enhancement post-transfer, trigger potent bystander activation of tumor-infiltrating host T cells.

Therefore, IL-36γ-induced Th9 cells demonstrate full effector function and super tumor-microenvironment immunomodulation that amplify endogenous T cell responses, establishing their exceptional therapeutic role in ACT that surpasses classic Th9 functional paradigms.

### 6. IL-36γ-programmed Th9 cells display less-exhausted and non-terminally differentiated T cell phenotype

Persistent antigen exposure in the tumor microenvironment promotes T cell terminal differentiation, apoptosis or exhaustion^9,42,43^. In view of the enhanced persistence of IL-36γ-programmed Th9 cells post-transfer, we hypothesized their resistance to exhaustion and/or delayed terminal differentiation compared to classic Th9 cells. TCellSI-based terminal exhaustion scoring confirmed significantly lower exhaustion scores in IL-36γ-programmed Th9 subsets versus classic counterparts (Figure 6A). GSEA further revealed attenuated expression of exhaustion-and terminal differentiation-related gene modules (GSE9650, GSE10239) in these subsets (Figure 6B and 6C). As key biomarkers of T cell exhaustion, immune checkpoints were analyzed. Heatmap demonstrated pronounced downregulation of *Pdcd1*, *Havcr2*, *Cd160*, *Tigit*, *Nt5e* and *Tox* alongside *Lag3* upregulation in IL-36γ-induced Th9 subsets compared to classic Th9 cells (Figure 6D). qRT-PCR corroborated these transcriptional patterns (Figure 6E). Flow cytometry showed substantial reductions in PD-1 and TIM-3 protein levels in both IL-36γ-induced Th9 cell subsets under anti-CD3/CD28 stimulation (Figure 6F and S6A), consistent with PD-1 downregulation observed under OVA-plused APC stimulation (Figure S6B). These data establish that IL-36γ-polarized Th9 cells retain exhaustion-resistant and non-terminally differentiated phenotypes *in vitro*.

**Figure 6.**
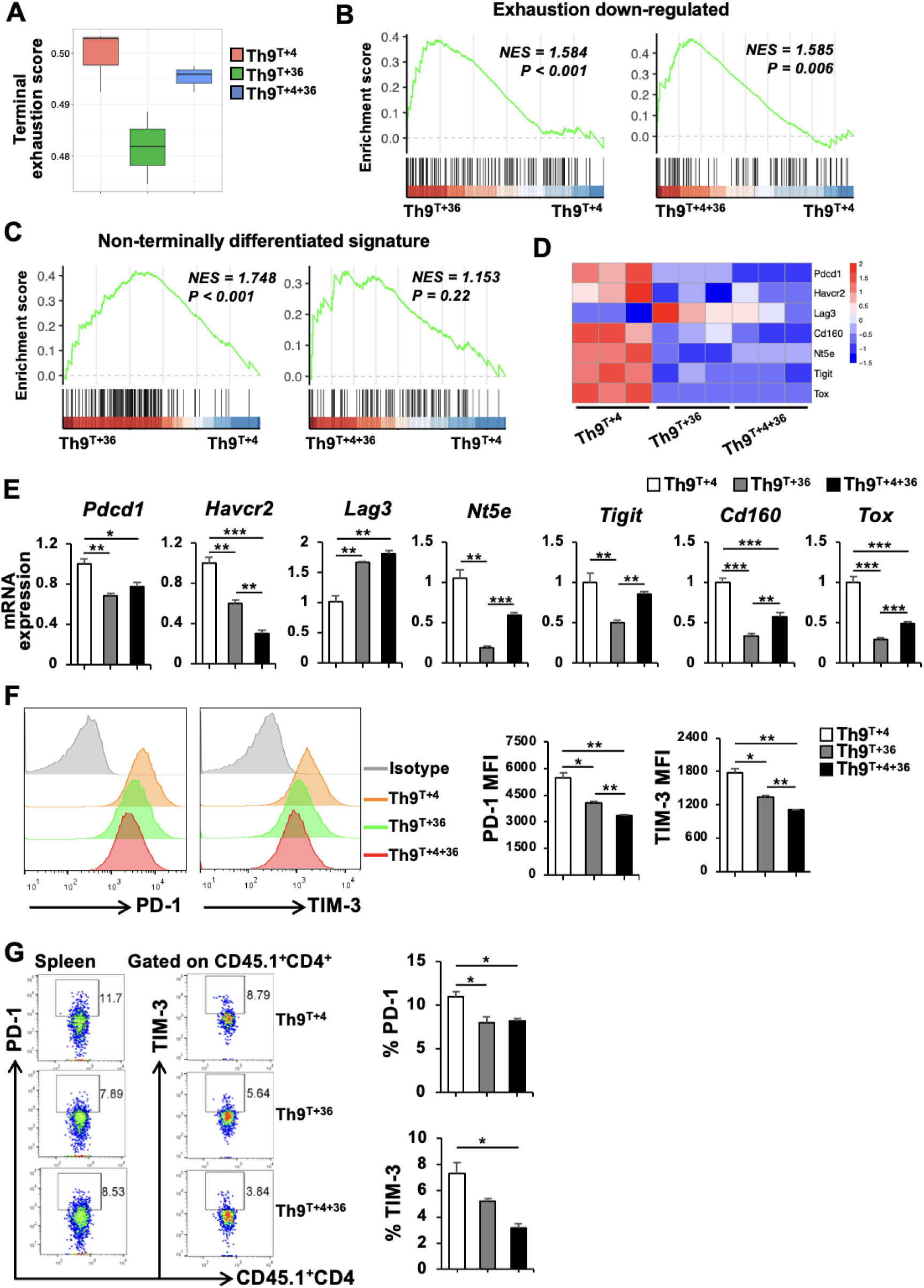
IL-36γ-programmed Th9 cells display less-exhausted and non-terminally differentiated T cell phenotypes. **(A)** Terminal exhaustion scores of Th9 subsets computed *via* TCellSI tool. **(B)** GSEA showing downregulation of exhaustion-associated genes (GSE9650) in IL-36γ-induced Th9 subsets. **(C)** Enrichment of non-terminally differentiated T cell signatures (GSE10239) across three Th9 subsets. **(D)** Heatmap demonstrating immune checkpoint molecules in Th9 subsets. **(E)** qRT-PCR validation of exhaustion-related transcripts in Th9 subsets. **(F)** Surface PD-1 and TIM-3 expression on *in vitro*-polarized Th9 subsets by flow cytometry (left, representative histograms; right, quantification). **(G)** Surface PD-1 and TIM-3 expression on adoptively transferred CD45.1^+^CD4^+^ T cells in spleens 10 days post-ACT (left, representative plots; right, quantification). One-way ANOVA was employed for data analysis **(E-G)**. \**p* < 0.05, \*\**p* < 0.01, \*\*\**p* < 0.001.

To evaluate *in vivo* consistence, we quantified PD-1 and TIM-3 on transferred CD45.1^+^CD4^+^ T cells in spleens, TdLNs and tumors 10 days post-adoptive transfer. IL-36γ-induced Th9 cell subsets displayed significantly reduced PD-1 and TIM-3 in spleens and PD-1 in TdLNs compared to classic Th9 cells, though TIM-3 in TdLNs, and PD-1 and TIM-3 in tumors remained comparable (Figure 6G, S6C and S6D). Collectively, IL-36γ-induced Th9 cells sustain reduced expression of checkpoint molecules *in vivo* compared to classic Th9 cells, preserving their exhaustion resistance and differentiation delay to prolong antitumor efficacy.

### 7. IL-36γ-programmed Th9 subsets exhibited stem-like and/or memory signatures

Stem-like memory T cells (Tscm), a self-renewing and long-lived population, sustain durable in *vivo* persistence. Given the superior longevity of IL-36γ-induced Th9 cells in tumor-bearing mice, we systematically evaluated their stem-like properties *via* TCellSI tool. Both IL-36γ-induced Th9 subsets displayed elevated stemness-associated gene scores versus classic Th9 cells (Figure 7A). Complementary GSEA revealed enriched central memory T cell (Tcm)-linked transcriptional programs (GSE26928) in IL-36γ-polarized Th9 cells (Figure 7B), suggesting their potential as a Tscm-and Tcm-like populations.

**Figure 7.**
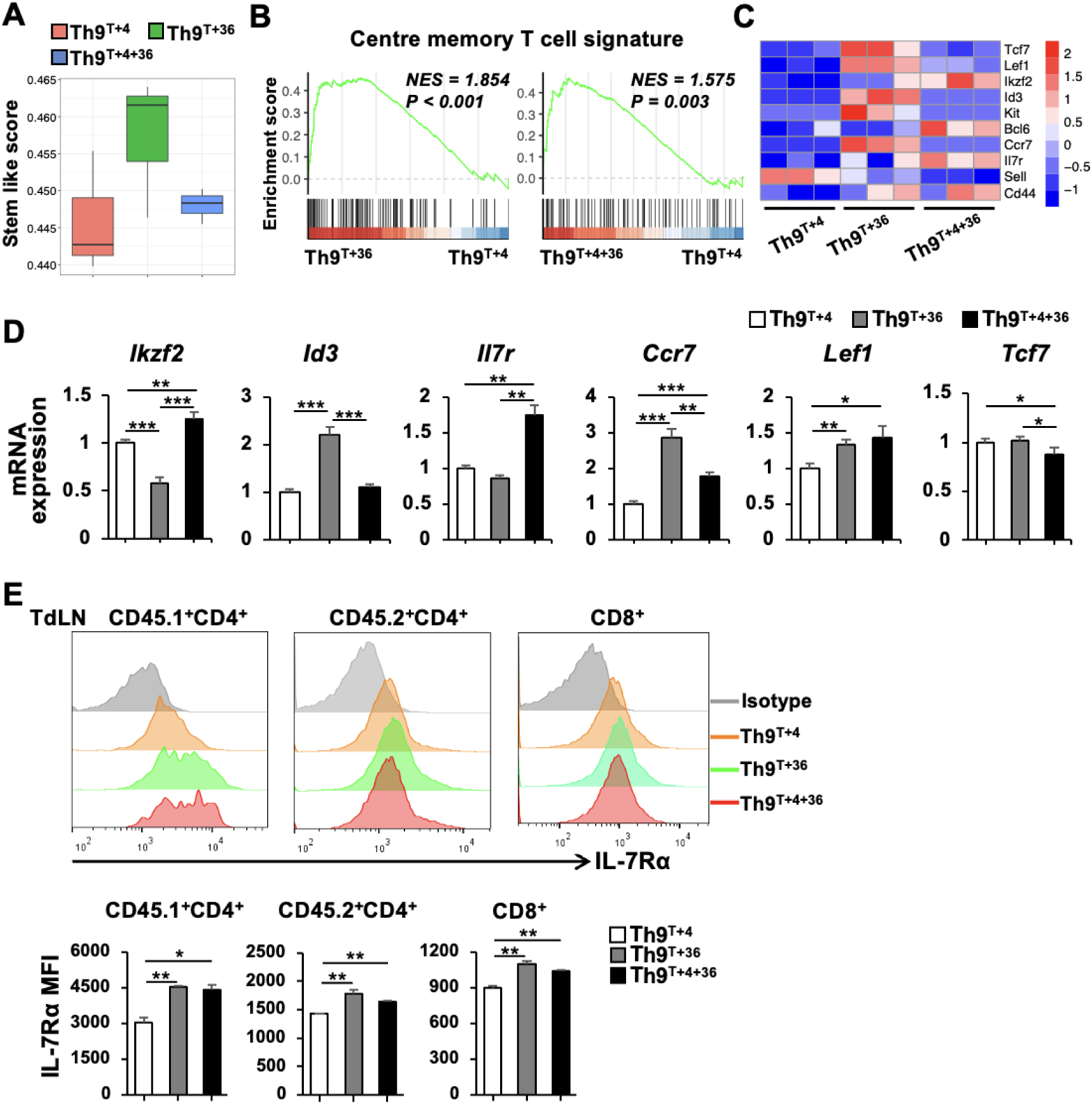
IL-36γ-programmed Th9 subsets exhibited stem-like and/or memory signatures. **(A)** Stemness scores of Th9 subsets calculated using TCellSI based on RNA-seq data. **(B)** GSEA showing enrichment of central memory T cell signatures (GSE26928) in IL-36γ-programmed Th9 subsets versus classic Th9 cells. **(C)** Heatmap of stemness-associated genes and memory markers across Th9 subsets. **(D)** qRT-PCR validation of stem/memory-related transcripts in IL-36γ-programmed Th9 subsets versus classic Th9 cells. **(E)** Flow cytometry qualified IL-7Rα expression on adoptively transferred CD45.1^+^CD4^+^ T, and endogenous CD45.2^+^CD4^+^ T and CD8^+^ T cells in TdLNs of tumor-bearing mice 20 days post-ACT across Th9 subset treatment groups (up, representative histograms; below, quantification). One-way ANOVA was employed for data analysis **(D and E)**. \**p* < 0.05, \*\**p* < 0.01, \*\*\**p* < 0.001.

Then, heatmap profiling of stemness and memory-associated genes identified IL-36γ-programmed Th9 cells possessed distinct transcriptional profiles. Compared to classic Th9 cells, Th9^TGF-β+IL-36γ^ subset exhibited upregulated transcripts of *Tcf7*, *Lef1, Id3*, *Kit and Ccr7*, while Th9^TGF-β+IL-4+IL-36γ^ subset demonstrated elevated expression of *Ikzf2, Bcl6 and Il7r* (Figure 7C). Meanwhile, qRT-PCR validation confirmed significant upregulation of *Lef1*, *Id3* and *Ccr7* in Th9^TGF-β+IL-36γ^ cells, and elevated expression of *Lef1*, *Ikzf2*, *Ccr7* and *Il7r* in Th9^TGF-β+IL-4+IL-36γ^ cells, compared to classic Th9 cells (Figure 7D).

Further, to functionally validate the stem-like and memory traits of IL-36γ-incduced Th9 subsets, we quantified IL-7Rα in adoptively transferred CD4^+^ T cells (CD45.1^+^) alongside host-derived CD4^+^ T and CD8^+^ T cells (CD45.2^+^), across spleen, TdLNs and tumor compartments. By day 20 post-transfer, the cohorts receiving IL-36γ-polarized Th9 subset therapies exhibited enhanced IL-7Rα expression compared to classic Th9 cells in transferred CD45.1^+^CD4^+^ T, and endogenous CD45.2^+^CD4^+^ T and CD45.2^+^CD8^+^ T cells in TdLNs but not in spleens and tumors (Figure 7E, S7A and S7B).

These findings collectively demonstrate that IL-36γ-polarized Th9 subsets *in vivo* acquire reinforced stem-like and/or memory phenotypes compared to classic Th9 cells, underpinned by conserved stemness modules and lymphoid niche retention after transfer.

## Discussion

While ACT has transformed cancer treatment, T cell exhaustion and poor persistence in tumors hinder its efficacy^42,44–46^. Th9/Tc9 cells offer distinct advantages over Th1/Tc1 subsets, characterized by IL-9 and IL-21 production and lower exhaustion-associated markers^10,27,47–49^. Preclinically, Th9/Tc9-polarized CAR-T cells outperform Th1/Tc1 counterparts through enhanced expansion, durable persistence, and superior tumor clearance^50,51^, establishing Th9/Tc9 programming as a transformative strategy to overcome ACT limitations in tumors. Notably, classic Th9 cells polarized with TGF-β and IL-4 exhibit suboptimal differentiation efficiency and their therapeutic potency can be further enhanced.

In the current study, we demonstrated that IL-36γ robustly drove Th9 generation. Furthermore, we identified a novel IκBζ-dependent mechanism through which IL-36γ conferred functional optimization to Th9 cells, marked by sustained persistence post-transfer, reduced exhaustion, enhanced stemness and memory properties, and potent immunomodulatory activity, positioning IL-36γ-induced Th9 cells as a superior ACT strategy.

Our findings demonstrated that IL-36γ induced Th9 cells at remarkable efficiencies, aligning with emerging strategies to augment Th9 differentiation *via* TNF-α signaling, yet distinctively operating *via* an alternatively IL-4-independent mechanism^21,28,29^. Importantly, IL-36γ-induced Th9 cells were identical to classic Th9 cells in cytokine profiles and transcription factors, yet exhibited amplified effector potential. Such stability contrasts with the reported capacity of IL-36γ to induce IL-9 in Th2 and iTreg subsets^33^, suggesting context-dependent regulation. This specificity may stem from IL-36γ’s unique engagement of IκBζ, which fine-tunes chromatin accessibility at *Il9* loci.

Mechanistically, IL-36γ drives Th9 cell differentiation dependent on enhancing STAT6, STAT5 and NF-κB pathways, the core regulators of classic Th9 differentiation^10,15,18,21,22,36,37^. STAT6, indispensable for IL-4-driven Th9 polarization^15,52,53^, proved equally critical for IL-36γ-induced Th9 cells, as STAT6 deficiency or inhibition abrogated IL-9 production. Similarly, IL-36γ augmented STAT5 signaling *via* IL-2, paralleling the reports that STAT5 sustains Th9 proliferation^54–56^. The non-canonical NF-κB pathway (RelB/p52) emerged as a key mediator, with IL-36γ upregulating *Nfkb2* (encoding p100/p52) and NF-κBp52 deficiency impairing IL-36γ-induced Th9 differentiation. This mirrors OX40-driven Th9 differentiation *via* RelB/p52 signaling pathway^29^. Such multipronged pathway activation ensures robust Th9 commitment, both in the presence and absence of IL-4.

A central breakthrough is the identification of IκBζ as a novel mediator governing Th9 differentiation. Notably, IκBζ has been identified as the key transcriptional regulator of IL-36 signaling in psoriasis^57^. While canonical and non-canonical NF-κB signaling pathways are well known to regulate Th9 cells^10,21,22,29,36,37^, our work revealed the specific requirement for the atypical nuclear IκB family member IκBζ in both IL-36γ-driven and classic Th9 polarization. Mechanistically, IκBζ directly bound to the *Il9* promoter and CNS-25 enhancer, the critical regulatory elements for Th9 lineage commitment, and *Nfkbiz* (encoding IκBζ) genetic ablation abolished IL-9 production even under IL-36γ stimulation. Therefore, IκBζ, alongside those conventional regulators such as PU.1, IRF4 and STAT6^15,16,58–60^, emerges as non-redundant transcriptional factor governing Th9 identity. Notably, this regulatory paradigm resembles the established role of IκBζ in regulating Th17 development, where it cooperates with RORγt/RORα to drive *Il17a* transcription^61^. The conserved requirement for IκBζ across these distinct T helper lineages suggests an evolutionary strategy where nuclear IκBζ license IL-9 or IL-17A production in conjunction with Th9 or Th17 lineage-specifying transcription factors.

Intriguingly, IκBζ exhibits context-dependent functional duality in T cell regulation. While essential for pro-inflammatory Th17 differentiation^61^, it simultaneously constrains Th1 polarization by suppressing IFN-γ production^62^ and regulates Treg functionality through TGF-β-dependent mechanisms^63^. Our findings that IκBζ drives Th9 differentiation extend this paradigm, revealing compartmentalized regulatory modules within the same molecule. The ability of IκBζ to interface with disparate signaling pathways underscores its role as a pleiotropic adaptor translating extracellular cues into T cell lineage-specific transcriptional programs.

IL-36γ-programmed Th9 cells eradicated established tumors more effectively than classic Th9 cells, attributable to their exceptional attributes as follows. Firstly, IL-36γ-polarized Th9 cells displayed the reduced expression of coinhibitory receptors such as PD-1 and TIM-3, contributing to evading terminal differentiation. Secondly, they possessed stem-like and memory traits resulting in lymphoid persistence. IL-36γ-driven Th9 cells exhibited elevated stemness scores and IL-7Rα expression. Upregulation of *Tcf7* and *Ccr7* further reflects a Tscm and Tcm phenotype, enabling lymphoid homing and self-renewal. Critically, these traits persisted *in vivo*, contrasting with Th1 cells that rapidly exhaust in TMEs. This durability mirrors CAR-T cells engineered with TCF7 overexpression^64^ but is achieved here through physiologic cytokine signaling. Thirdly, IL-36γ endowed Th9 cells with potent bystander activation of endogenous immunity. While IL-36γ-induced Th9 subsets *in vivo* showed comparable intrinsic cytotoxicity to classic Th9 cells post-transfer, they potently activated endogenous CD4^+^ T and CD8^+^ T cells. This aligns with the reports that classic Th9 cells remodel tumor microenvironments^27,48^ but IL-36γ-programmed Th9 cells may enhance this crosstalk. Mechanistically, IL-36γ-driven *Il21* transcription likely underpins this effect, as IL-21 is critical for sustaining antitumor immunity^65,66^.

Current ACT platforms, including CAR-T cells, face hurdles in solid tumors due to T cell exhaustion and poor infiltration. IL-36γ-induced Th9 cells address these limitations and remodel the tumor microenvironments. Their stem-like and memory properties suggest potential for long-term immunosurveillance, critical for preventing relapse.

This study establishes several research trajectories. Firstly, elucidating the interplay of IκBζ with other Th9 regulators can refine differentiation protocols. Secondly, engineering CAR-T cells with IL-36γ or IκBζ overexpression may enhance their efficacy. Thirdly, combining IL-36γ-induced Th9 cells with PD-1 blockade may synergistically amplify responses.

In conclusion, IL-36γ-induced Th9 cells represent a paradigm shift in ACT, merging sustained persistence, cytotoxicity, and microenvironmental reprogramming. By unraveling the IκBζ-dependent mechanism, this work not only advances Th9 biology but also charts a path toward curative immunotherapies for refractory cancers.

## Methods and Methods

### Animal experiments

All animal experiments were performed using 6–8-week-old C57BL/6j female mice with strict age matching within experimental groups. Procedures were conducted in a specific pathogen-free (SPF) facility approved by the Soochow University Animal Ethics Committee (Approval ID: 202010A624). Mice were housed under standardized conditions with a 12:12-hour light-dark cycle in open-top cages, provided ad libitum access to food and water. Health monitoring was performed routinely, with intensified surveillance when anticipating adverse effects. The following mouse strains were utilized: CD45.2, CD45.1, *Cd4*^cre^*Nfkb2*^fl/fl^ and *Cd4*^cre^*Nfkbiz*^fl/fl^ strains (purchased from Jiangsu GemPharmatech Co., Ltd, China); *Stat6*^-/-^ mice (kindly provided by Prof. Y. Qin) and CD45.1 OT-II mice (generated through crossing CD45.1 with OT-II strains).

### Cell line

The B16-OVA melanoma cell line, generously provided by Prof. B. Lu, was cultured in RPMI 1640 medium (Hyclone) supplemented with 10% heat-inactivated fetal bovine serum (Gibco), 100 U/mL penicillin and 100 μg/mL streptomycin (Invitrogen) under standard humidified incubator conditions (37°C, 5% CO_2_).

### Reagents

Recombinant murine TGF-β, IL-4, IL-36γ, IL-2, IL-12 and GM-CSF were purchased from R&D systems. The neutralizing mAbs anti-mouse IFN-γ (clone XMG1.2) and anti-mouse IL-4 (clone 11B11) were sourced from BioXCell. Biotin-conjugated anti-mouse CD4 mAb (clone GK1.5), purified anti-mouse CD3ε mAb (clone 145-2C11) and anti-mouse CD28 mAb (clone 37.51), Fixation/Permeabilization Buffer, Intracellular Perm Wash Buffer, True-Nuclear™ Fixation Buffer and True-Nuclear™ Perm Buffer were acquired from BioLegend. Streptavidin MicroBeads was obtained from Miltenyi Biotec. STAT6 inhibitor AS1517499, NF-κB inhibitor QNZ and mitomycin C were obtained from Selleck Chemicals. STAT5 inhibitor 573108-M was purchased from Millipore. Cyclophosphamide (CTX) and phorbol 12-myristate 13-acetate (PMA) were purchased from Sigma-Aldrich. Ionomycin calcium salt and brefeldin A (BFA) were acquired from BioGems and Tocris Bioscience, respectively. ACK lysing buffer was purchased from Thermo Fisher Scientific. OVA_323-339_ peptide (ISQAVHAAHAEINEAGR-NH_2_) was ordered from Abgent.

### *In vitro* Th subset differentiation

Spleens and lymph nodes were isolated from CD45.2, CD45.1 OT-II, *Stat6*^-/-^, *Cd4*^cre^*Nfkb2*^fl/fl^ or *Cd4*^cre^*Nfkbiz*^fl/fl^ mice and then single-cell suspensions were prepared by mechanical dissociation followed by erythrocyte lysis using ACK lysing buffer. Subsequently, naïve CD4^+^ T cells were purified by magnetic bead-based methods. For APC preparation, splenocytes underwent negative selection using biotin-anti-mouse CD3ε mAb. APCs were plated at a density of 2 × 10^6^ cells/mL per well in 6-well plates, pulsed with 5 μg/mL OVA_323-339_ peptide and treated with 40 μg/mL mitomycin C to generate OVA323-339-loaded APCs.

Purified naïve CD4^+^ T cells were activated either with plate-bound anti-mouse CD3ε mAb (5 μg/mL) combined with soluble anti-mouse CD28 mAb (5 μg/mL) or through co-culture with OVA323-339-loaded APCs. For Th subset polarization, distinct cytokine cocktails were supplemented as follows. Th1 cells: IL-12 (10 ng/mL), IL-2 (20 ng/mL) and anti-IL-4 mAb (20 μg/mL); classic Th9 cells: TGF-β (2.5 ng/mL), IL-4 (10 ng/mL), IL-2 (20 ng/mL), anti-IFN-γ mAb (20 μg/mL); Th9^TGF-β+IL-36γ^ cells: TGF-β (2.5 ng/mL), IL-36γ (100 ng/mL), IL-2 (20 ng/mL), anti-IFN-γ mAb (20 μg/mL); Th9^TGF-β+IL-4+IL-36γ^ cells: TGF-β (2.5 ng/mL), IL-4 (10 ng/mL), IL-36γ (100 ng/mL), IL-2 (20 ng/mL), anti-IFN-γ mAb (20 μg/mL). In some specific experiments, IL-36γ was added at a concentration gradient (0, 1.5625, 3.125, 6.25, 12.5, 25, 50 and 100 ng/ml). All cells were cultured in RPMI 1640 medium supplemented with 10% fetal bovine serum, 100 U/mL penicillin and 100 μg/mL streptomycin under standard humidified incubator conditions (37°C, 5% CO_2_). After 2 days or 36 h of culture, cells and supernatants were harvested for flow cytometry, qRT-PCR, ELISA or RNA-SEQ analyses. In some experiments, STAT6 inhibitor (AS1517499), STAT5 inhibitor (573108-M) or NF-κB inhibitor (QNZ) were added with indicated concentrations during culture initiation.

### Flow cytometry

*In vitro* differentiated Th9 cells or cells isolated from mouse spleens, TdLNs and tumors were first stained with eBioscience™ Fixable Viability Dye (Thermo Fisher Scientific) for 30 min in 4 °C, then underwent surface staining for 20 min. For intracellular cytokine analysis, cells were activated with a PMA/ionomycin cocktail (500 ng/mL PMA, 1 μg/mL ionomycin) for 1 h following treatment with BFA (10 μg/mL) for 3 h, then cells were fixed and permeabilized post eBioscience™ Fixable Viability Dye and surface staining. Subsequently, permeabilized cells were stained with cytokine antibodies for 30 min in 4°C prior to flow cytometry analysis. For nuclear Ki67 determination, True-Nuclear™ Fixation Buffer and True-Nuclear™ Perm Buffer were used. All fluorochrome-conjugated antibodies were sourced from BioLegend as follows: APC-conjugated anti-mouse CD45 (clone 30-F11), APC/Cy7-conjugated anti-mouse CD45.1 (clone A20), Pacific Blue-conjugated anti-mouse CD8α (clone 53-6.7), BV605-conjugated anti-mouse CD4 (clone GK1.5), PE-conjugated anti-mouse PD-1 (clone 29F.1A12), PerCP/Cy5.5-conjugated anti-mouse TIM-3 (clone B8.2C12), FITC-conjugated anti-mouse IFN-γ (clone XMG1.2), PerCP/Cy5.5-conjugated anti-mouse TNF-α (clone MP6-XT22), FITC-conjugated anti-mouse Ki67 (clone 16A8), PerCP/Cy5.5-conjugated anti-mouse IL-7Rα (clone A7R34), and eFluor 660-conjugated anti-mouse IL-9 (clone RM9A4). Fixable Viability Dye eFluor™ 780 was obtained from Thermo Fisher Scientific.Flow cytometry data were acquired on the CytoFLEX platform and analyzed using FlowJo software (Treestar).

### ELISA

Concentrations of IL-9 and IL-10 in the culture supernatant were determined by ELISA kits (Thermo Fisher Scientific), according to the manufacturers’ instructions.

### shRNA knockdown

ShRNA knockdown lentiviral particles targeting mouse *Batf3* (sequence: 5’-GAAAGTTCGAAGGAGAGAGAA-3’) and non-targeting control shRNA (sequence: 5’-TTCTCCGAACGTGTCACGT-3’) were synthesized by GENECHEM (Shanghai, China) using the GV644 lentiviral vector (pRRLSIN-cPPT-U6-shRNA-SFFV-EGFP-SV40-puromycin). Naïve CD4⁺ T cells isolated from mouse spleens were cultured under different Th9-polarizing conditions. At 24 hours post-culture, cells were transduced with *Batf3*-targeting or control shRNA lentiviral particles at a multiplicity of infection (MOI) of 40. After 12-hour incubation, the supernatant was replaced with fresh different Th9-polarizing medium. Cells were harvested 30 hours later for flow cytometry analysis of intracellular IL-9 and ELISA quantification of IL-9 levels in culture supernatants.

### Tumor model and adoptive cell therapy

For generation of therapeutic antigen-specific Th subset cells, CD4^+^ T cells purified from CD45.1 OT-II mice were co-cultured with OVA_323-339_-loaded APCs (CD45.2^+^) at 3 × 10^5^: 9 ×10^5^ ratio in 48-well plates under Th1, Th9^TGF-β+IL-4^, Th9^TGF-β+IL-36γ^ or Th9^TGF-β+IL-4+IL-36γ^ polarized conditions. For preparation of BMDCs, bone marrow cells isolated from femurs and tibiae of CD45.2 mice were filtered through 70-μm strainers, lysed with ACK buffer, and cultured at 6 × 10^6^ cells in 6-well plates in RPMI-1640 medium with 10% heat-inactivated fetal bovine serum, supplemented with GM-CSF (20 ng/mL) and IL-4 (20 ng/mL). On day 6, cells were pulsed with 5 μg/mL OVA_323-339_ peptide and then mature BMDCs were harvested on day 7 for adoptive transfer.

A B16-OVA melanoma model was established by subcutaneous injection of 2×10⁵ tumor cells into the flanks of CD45.2 mice. On day 5 post-implantation, mice received CTX (200 mg/kg) or PBS preconditioning. On day 6, CTX-treated cohorts were administered 4×10^5^ OVA_323-339_-pulsed BMDCs with 2 × 10^6^ antigen-specific CD45.1^+^CD4^+^ T subsets (Th1, Th9^TGF-β+IL-4^, Th9^TGF-β+IL-36γ^, Th9^TGF-β+IL-4+IL-36γ^). Tumor size was measured every 2 days. Mice were euthanized on day 10 and 20 post-treatment for immune profiling. Single-cell suspensions from spleens, TdLNs and tumor tissues were prepared through mechanical dissociation and filtration for flow cytometry analysis.

### RNA extraction and qPCR with reverse transcription

Total RNA was extracted using the RNeasy Plus Micro Kit (QIAGEN), followed by cDNA synthesis with the High-Capacity cDNA Reverse Transcription Kit (Applied Biosystems). Amplification reactions were carried out on a StepOne Plus Real-Time PCR System (Applied Biosystems) using SYBR Green Master Mix (Applied Biosystems). Primer sequences are listed in Supplemental information Table 1. Gene expression was normalized to the expression of the housekeeping gene β-actin.

### RNA sequencing and analysis

Total RNA was extracted as above. RNA purity and quantification were evaluated using the NanoDrop 2000 spectrophotometer (Thermo Fisher Scientific). RNA integrity was assessed using the Agilent 2100 Bioanalyzer (Agilent Technologies). Then the libraries were constructed using VAHTS Universal V6 RNA-seq Library Prep Kit according to the manufacturer’s instructions. The transcriptome sequencing and analysis were conducted by OE Biotech. The libraries were sequenced on a llumina Novaseq 6000 platform and 150 bp paired-end reads were generated. Raw reads of fastq format were firstly processed using fastp and the low-quality reads were removed to obtain the clean reads^67^. The clean reads were mapped to the reference genome using HISAT2^68^. FPKM^69^ of each gene was calculated and the read counts of each gene were obtained by HTSeq-count^70^.

PCA analysis were performed using R (v4.4.1) to evaluate the biological duplication of samples. Differentially expressed genes (adjusted *p*-value < 0.01, LogFC > 0.5) were determined by DESeq2 (v1.46.0)^71^.

Venn diagrams were plotted with the R package venn (v1.12) and VennDiagram (v1.7.3). Heat maps were plotted with the pheatmap (v1.0.12). TCellSI analysis was performed with the TCSS_Caluclate functions using TCellSI package (v0.1.0) with default parameters^41^.

### Gene-set and pathway enrichment analysis

GSEA was done with the GSEA functions from clusterProfiler package (v4.14.3) using default parameters^72^. Gene ontology enrichment analysis (GO Biological Processes) was performed with the enrichGO functions using clusterProfiler with default parameters^73^. Multiple comparative analysis was performed using datasets publicly available through the Gene Expression Omnibus (GEO) database under the following accession numbers: GSE222909 (Th9 signature), GSE22886 (Th1 signature), GSE41867 (Effector T cell signature), GSE9650 (Exhaustion down-regulated), GSE10239 (non-terminally differentiated signature), GSE26928 (Centre memory T cell signature). A list of public datasets used can be found in Supplemental information Table 2.

### ChIP-qPCR assay

Chromatin crosslinking, nuclei isolation and chromatin fragmentation were performed with Magna ChIP Kit (Merck). Anti-IκBζ antibody (Cell Signaling Technologies, 1:50) and Normal Mouse IgG antibody (Merck, 1:500) were used to pull down sheared chromatin from CD4^+^ T cells. For ChIP-qPCR analysis, the following DNA fragments were amplified with primers as described: *Il9* promoter F: 5′-GTGGGCACTGGGTATCAGTTTGATGT-3′, *Il9* promoter R: 5′-CAGTCTACCAGCATCTTCCAGTCTAG-3′; *Il9* CNS25 F: 5′-AGCAGGCGACCACTTTAAAA-3′, *Il9* CNS25 R: 5′-GCCAACTCTCAGCATGTGTT-3′.

### Statistical analysis

All *in vitro* experiments comprised 3 biological replicates and *in vivo* studies included at least 2 independent trials, each containing ≥ 5 mice per group. Data presented as mean ± s.e.m. Statistical analyses were performed using GraphPad Prism v10.0. Unpaired Student’s *t*-test, one-and two-way ANOVA were used to analyze the comparisons between groups. Differences between survival curves were analyzed using the log-rank test. All statistical tests were two-sided. \**p* < 0.05, \*\**p* < 0.01, \*\*\**p* < 0.001.

## DATA AVAILABILITY

Our RNA-seq data generated in this study are deposited in the National Center for Biotechnology Information Gene Expression Omnibus under accession number GSE318950. Publicly available files corresponding to the GEO accession codes: GSE222909, GSE22886, GSE41867, GSE9650, GSE10239 and GSE26928.

## ACKNOWLEDGMENTS

This work was funded by National Natural Science Foundation of China (NSFC) grants (31970833, 91642103, 82073180, 81802572), a project funded by the Priority Academic Program Development of Jiangsu Higher Education Institutions, Jiangsu Provincial Natural Science Foundation Project (BK20231201), the Project of State Key Laboratory of Radiation Medicine and Protection, Soochow University (GZK1202305, GZK1202203), the Suzhou City Key Clinical Disease Diagnosis and Treatment Technology Special Project (LCZX202216), the Science and Technology Development Plan of Suzhou Science and Technology Bureau (SYW2024013), the Suzhou City health Commission project (DZXYJ202411), the Key Project of Science and Education Promoting Health of Suzhou City (ZDXM2024003), the Research of Medical Innovation and Application, Science and Technology Development Project of Suzhou City (SKY2023109), the Gusu Talent Program of Suzhou City (GSWS2024023). All the funders were not involved in any aspect of the study design, data handling, manuscript preparation or publication decision.

## AUTHOR CONTRIBUTIONS

Q. Z., Y. C., J. Z. and X. W. conceived and designed the study. Q. Z., Y. C., J. G., P. T., H. J., Z. M., X. L. and X. J. performed the experiments and analyzed the data. Z. H. and Y. Z. assisted with animal experiments and data analysis. Q. Z. and Y. C. wrote the original draft of the manuscript. X. Z., J. Z. and X. W. supervised the project, interpreted results and revised the manuscript critically. All authors read and approved the final manuscript.

## Supplemental materials

**Figure S1.**
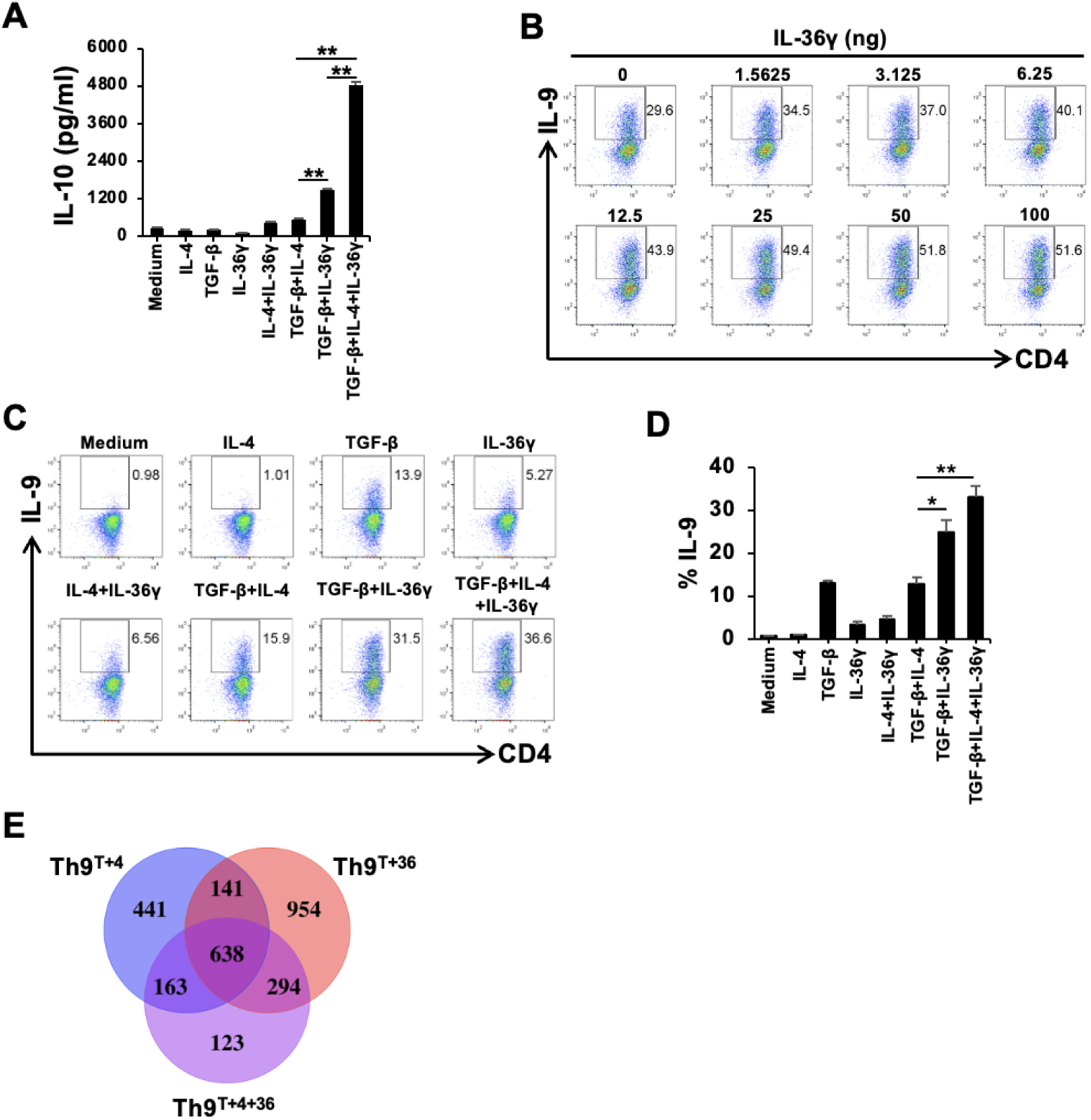
IL-36γ promotes the generation of Th9 cells with transcriptomic profiles comparable to those of classic Th9 cells. **(A)** Naïve CD4^+^ T cells from OT-II mice were activated with OVA_323-339_ peptide-pulsed APCs and cultured in the presence of TGF-β (2.5 ng/ml), IL-4 (10 ng/ml), IL-36γ (100 ng/ml) or in indicated combinations for 2 days. IL-10 levels in supernatants were quantified by ELISA. **(B)** Naïve CD4^+^ T cells from OT-II mice were activated with OVA_323-339_-pulsed APCs and cultured under classic Th9-polarizing conditions with graded doses of IL-36γ as indicated for 2 days. Intracellular IL-9 expression in CD4^+^ T cells was analyzed by flow cytometry. Data were representative of one experiment out of two independent replicates. **(C and D)** Naïve CD4^+^ T cells were stimulated with plate-bound anti-CD3/CD28 mAbs (5 μg/ml each) and IL-2 (20 ng/ml) and cultured in the presence of TGF-β (2.5 ng/ml), IL-4 (10 ng/ml), IL-36γ (100 ng/ml) or in indicated combinations for 2 days. **(C)** Representative flow cytometry plots of IL-9^+^ cells. **(D)** Quantitative analysis of IL-9^+^ cell frequencies. **(E),** Venn diagram analysis demonstrating shared transcriptional signatures among Th9^T+4^, Th9^T+36^ and Th9^T+4+36^ cell subsets. For all *in vitro* experiments, one-way ANOVA was used **(A and D)**. All statistical tests were two-sided and all replicates were biologically independent samples (*n* = 3). \**p* < 0.05, \*\**p* < 0.01, \*\*\**p* < 0.001.

**Figure S2.**
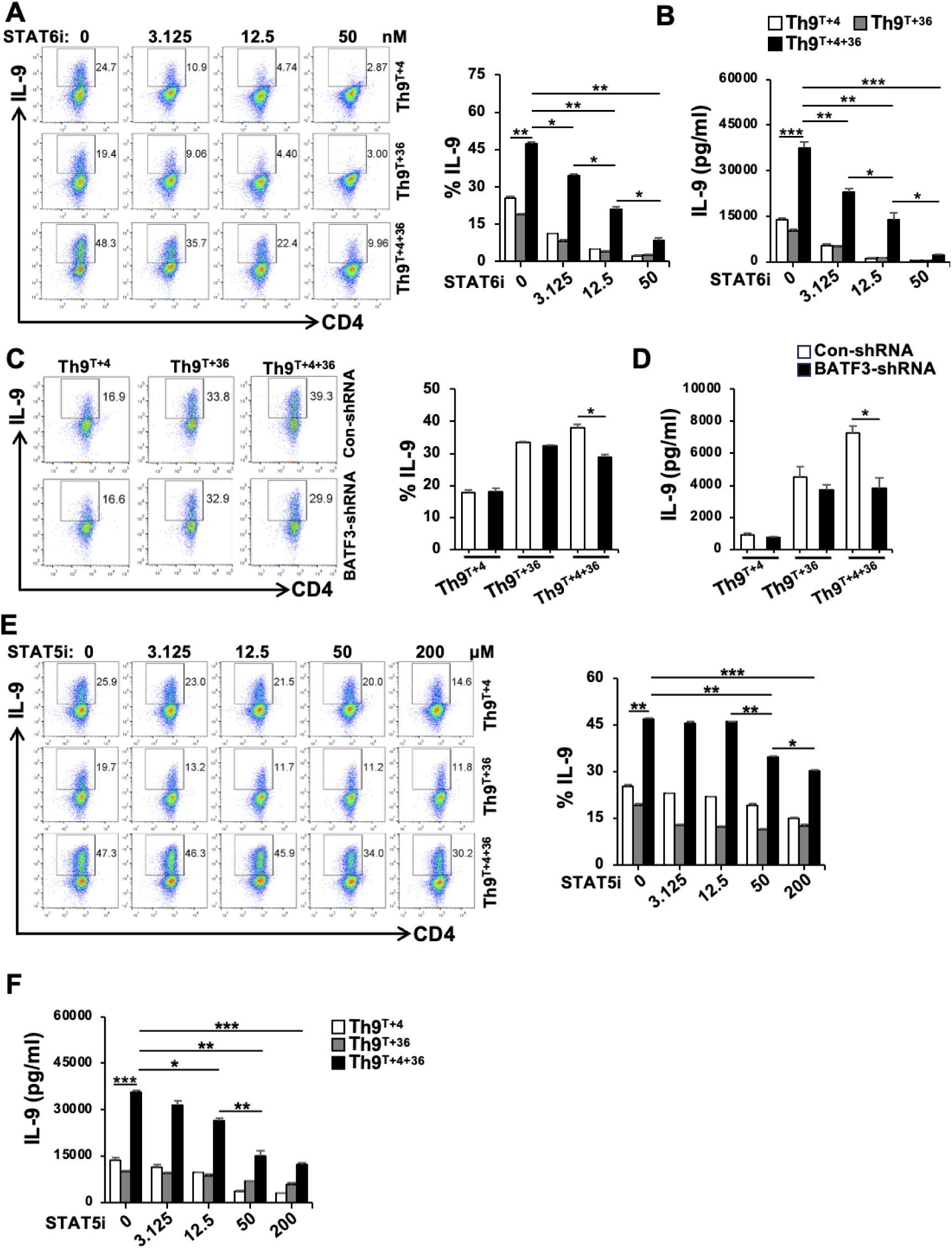
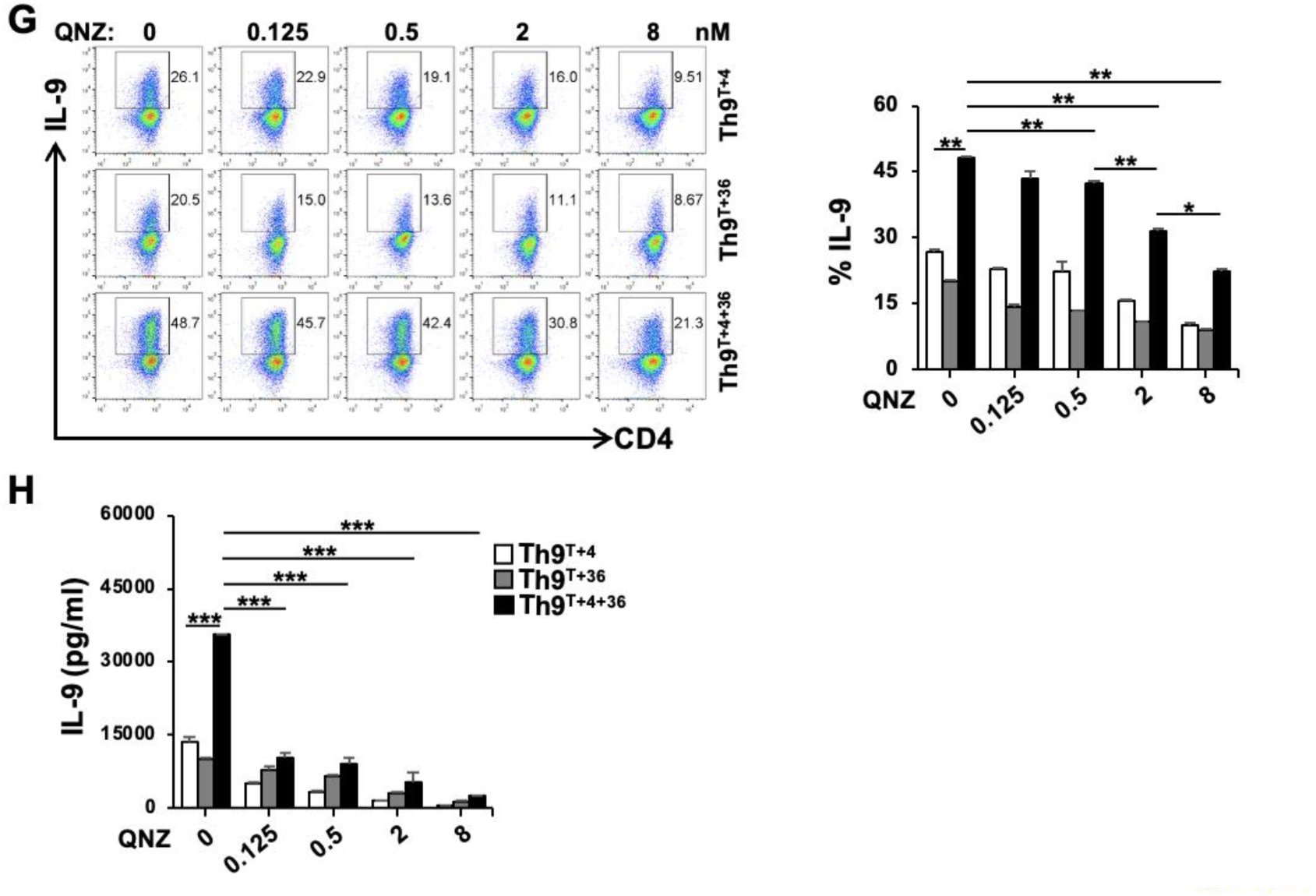
IL-36γ drives Th9 cell differentiation dependent on STAT6, BATF3, STAT5 and NF-κB signaling pathways. (A and. **B)** Naïve CD4^+^ T cells from OT-II mice were activated with OVA_323-339_-pulsed APCs and cultured under Th9^TGF-β+IL-4^, Th9^TGF-β+IL-36γ^ or Th9^TGF-β+IL-4+IL-36γ^ conditions with graded concentrations of STAT6 inhibitor (AS1517499) as indicated for 2 days. **(A)** Flow cytometric analysis of representative plots (left) and quantification (right) for IL-9^+^ cells. **(B)** IL-9 secretion in supernatants measured by ELISA. **(C and D)** Naïve CD4^+^ T cells were activated with plated-bound anti-CD3/CD28 mAbs (each 5 μg/ml) and IL-2 (20 ng/ml) under Th9^TGF-β+IL-4^, Th9^TGF-β+IL-36γ^ or Th9^TGF-β+IL-4+IL-36γ^ conditions and transduced with control *shRNA* or *Batf3*-targeting *shRNA* prior to cell polarization. **c,** Flow cytometric analysis of representative plots (left) and quantification (right) for IL-9^+^ cells. **d,** Supernatant IL-9 levels measured by ELISA. **(E and F)** Naïve CD4^+^ T cells from OT-II mice were cultured under conditions as **(A and B)** with increasing concentrations of STAT5 inhibitor (573108-M). Cells and supernatants were harvested after 2 days. **(E)** Flow cytometric analysis of representative plots (left) and quantification (right) for IL-9^+^ cells. **(F)** IL-9 levels in culture supernatants measured by ELISA. **(G and H)** Naïve CD4^+^ T cells from OT-II mice were activated and cultured under conditions as **(A and B)** with escalating doses of the NF-κB inhibitor QNZ as indicated. Cells and supernatants were harvested after 2 days. **(G)** Flow cytometric analysis of representative plots (left) and quantification (right) for IL-9^+^ cells. **(H)** IL-9 levels in culture supernatants measured by ELISA. A two-way ANOVA was used for comparisons of dose-dependent studies with three groups and multiple dose points **(A, B and E-H**. Comparisons between Control and *Batf3* shRNA groups within the same Th9 subset were performed using unpaired Student’s *t*-test **(C and D)**. All statistical tests were two-sided and all replicates were biologically independent samples (*n* = 3). \**p* < 0.05, \*\**p* < 0.01, \*\*\**p* < 0.001.

**Figure S3.**
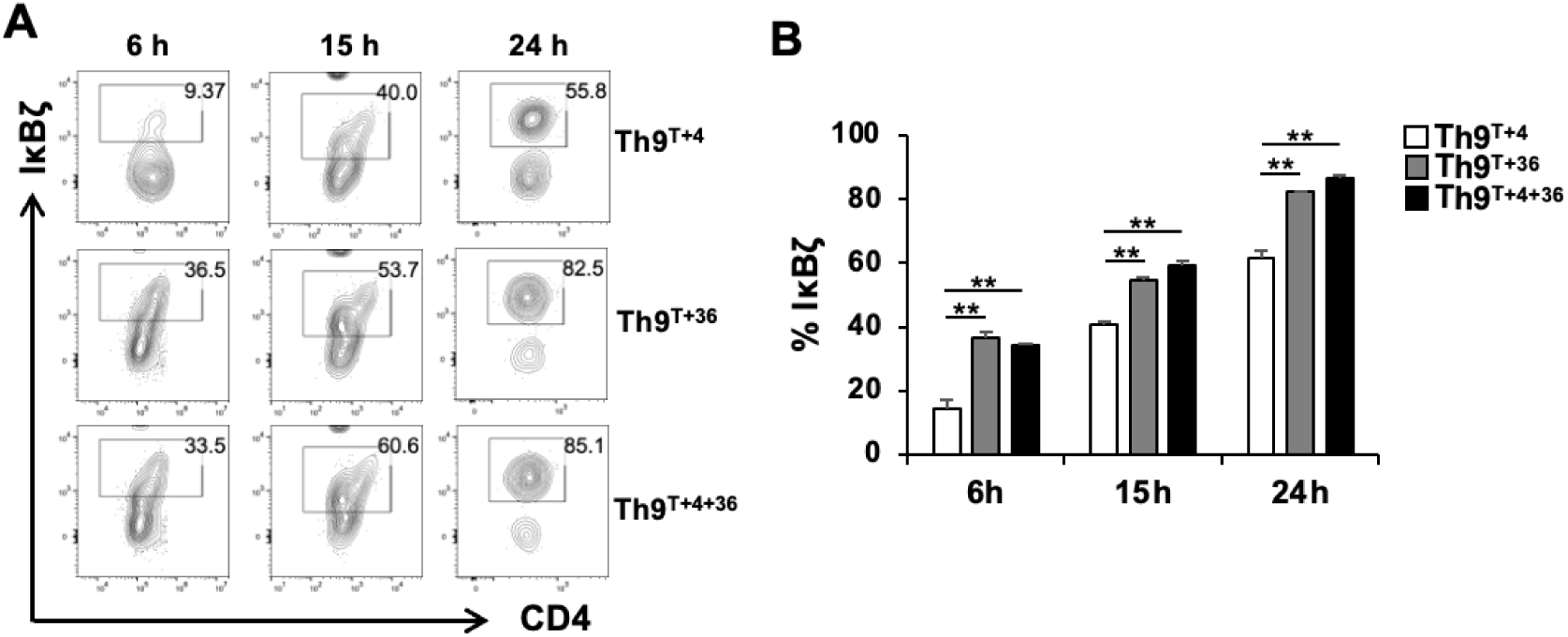
IκBζ exhibits enhanced nuclear accumulation in Th9^TGF-β+IL-36γ^ and Th9^TGF-β+IL-4+IL-36γ^ subsets compared to classic Th9 cells. (A-C) Naïve CD4^+^ T cells were activated with anti-CD3/CD28 mAbs (each 5 μg/ml) and IL-2 (each 20 ng/ml) and cultured under different Th9 subset-polarizing conditions as indicated. Cells were harvested at 6, 15 and 24 h. **(A and B)** Time-dependent IκBζ levels accumulated in the nucleus were quantified by flow cytometry. **(A)** Representative flow cytometry plots. **(B)** Quantitative analysis of IκBζ in the nucleus across Th9 subsets. This result is representative of one of two independent experiments. One-way ANOVA was employed to compare differences among the three Th9 subsets at identical time points **(B)**. All replicates were biologically independent samples (*n* = 3). \*\**p* < 0.01.

**Figure S4.**
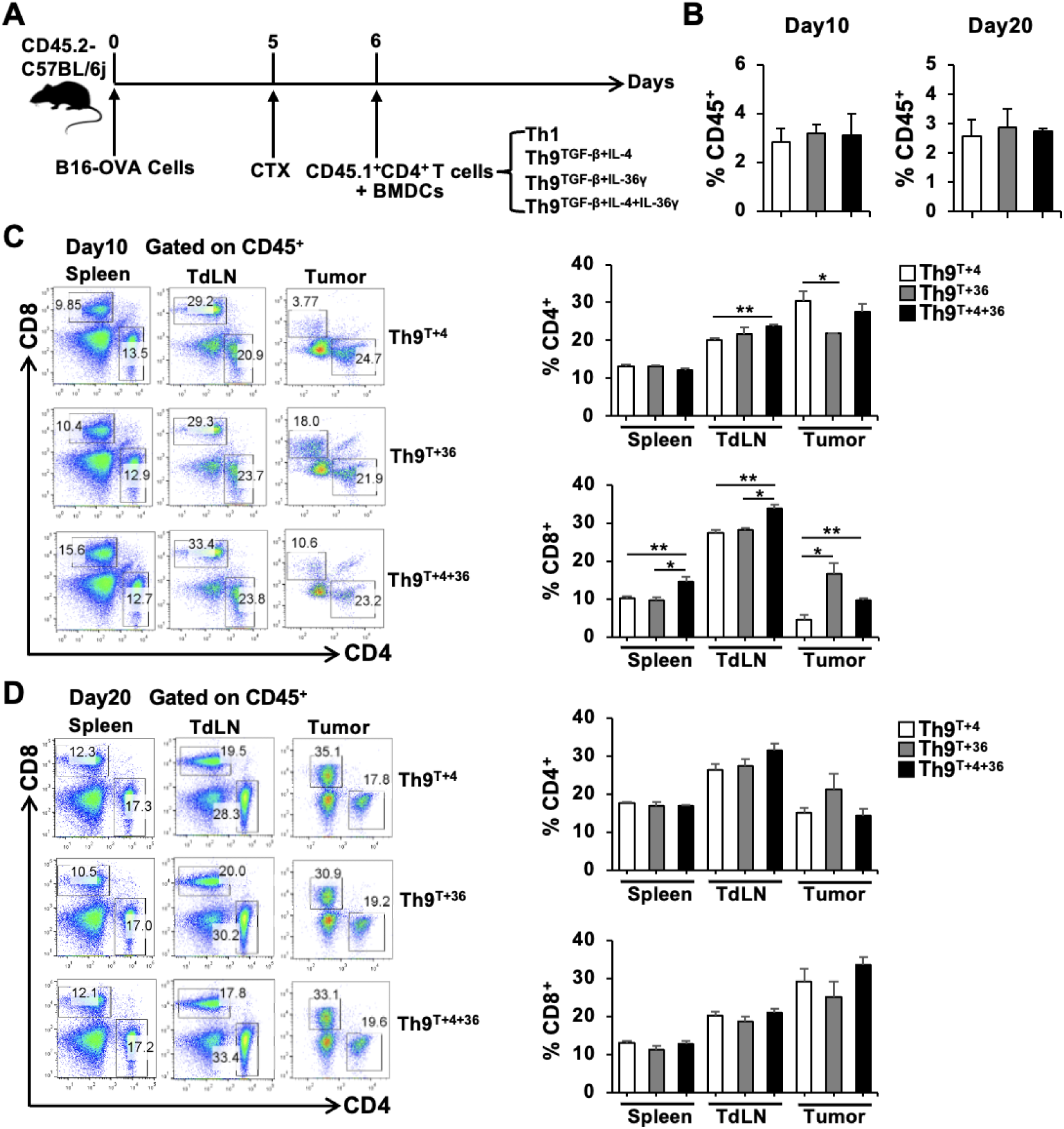
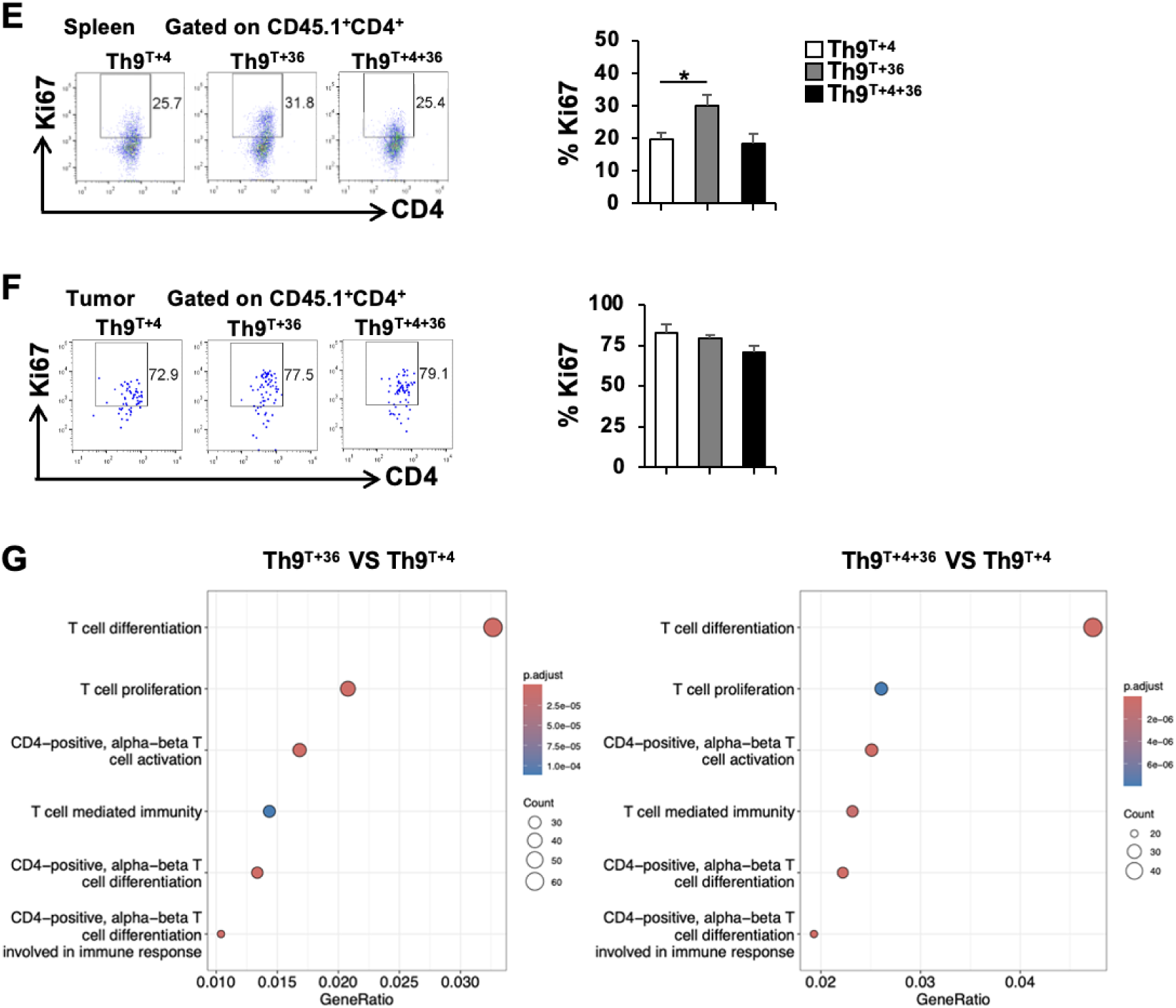
IL-36γ-programmed Th9 cells potentiate ACT through expanding endogenous T cells and possess the features of enhanced proliferation, activation and lineage commitment. **(A)** B16-OVA melanoma-bearing mice (CD45.2^+^) (*n* = 5) received CTX or PBS pretreatment on day 5 post-tumor inoculation, followed by adoptive transfer of OVA-specific CD45.1^+^Th1, CD45.1^+^Th9^TGF-β+IL-4^, CD45.1^+^Th9^TGF-β+IL-36γ^ or CD45.1^+^Th9^TGF-β+IL-4+IL-36γ^ subsets, respectively combined with OVA_323-339_-pulsed BMDCs on day 6. Tumor volumes were monitored every 2 days. **(B-F)** Mice were sacrificed on day 10 and 20 post-treatment for immune profiling. **(B)** The frequency of total CD45^+^ leukocyte infiltration in tumors across differenTh9 subset treatment groups. **(C and D)** Frequencies of CD4^+^ T and CD8^+^ T cells within CD45^+^ populations across spleens, TdLNs and tumors on day 10 **(C)** and day 20 **(D)** (left, representative flow plots; right, quantification) after therapy. **(E and F)** Flow cytometric analysis of representative plots (left) and quantification (right) for Ki67^+^ proliferation rates in spleen-**(E)** and tumor-**(F)** resident adoptively transferred CD45.1^+^CD4^+^ T cells on day 10. **(G)** GO enrichment analysis of IL-36γ-programmed Th9 subsets compared to classic Th9 cells. All quantified data **(B-F)** were analyzed for intergroup differences using one-way ANOVA across three Th9 therapeutic cohorts in the spleens, TdLNs and tumor tissues. \**p* < 0.05, \*\**p* < 0.01.

**Figure S5.**
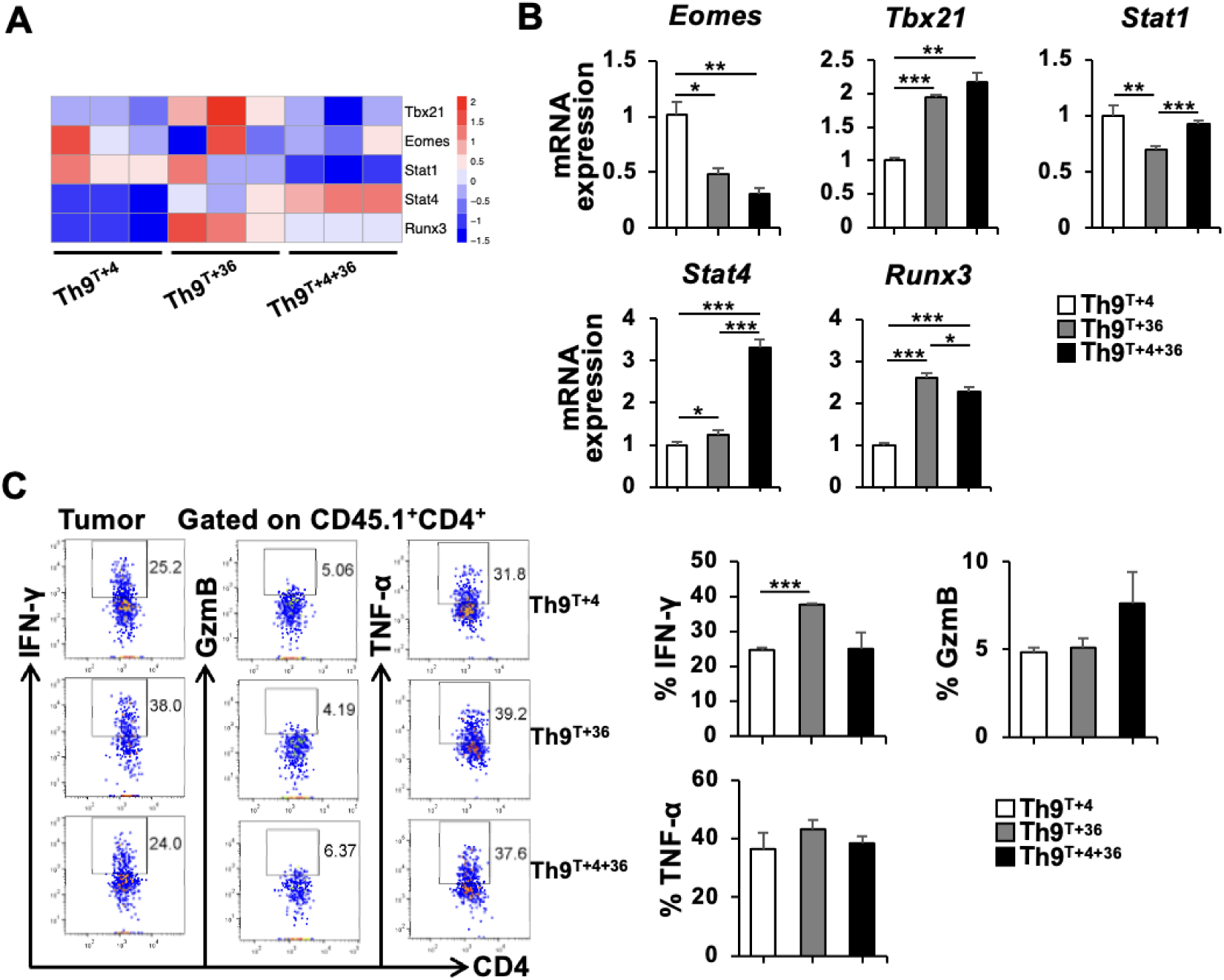
IL-36γ-induced Th9 cells display distinct Th1-like transcriptional programs and maintain full effector functions. **(A)** Heatmap showing expression levels of Th1-associated transcription factors in three Th9 subsets. **(B)** qRT-PCR validation of Th1-associated transcription factors across three Th9 subsets. **(C)** Flow cytometric analysis of the expression of IFN-γ, GzmB and TNF-α in CD45.1^+^CD4^+^ T cells in tumors 10 days post-adoptive transfer (left, representative plots; right, quantification). One-way ANOVA was employed for data analysis **(B and C)**. \**p* < 0.05, \*\**p* < 0.01, \*\*\**p* < 0.001.

**Figure S6.**
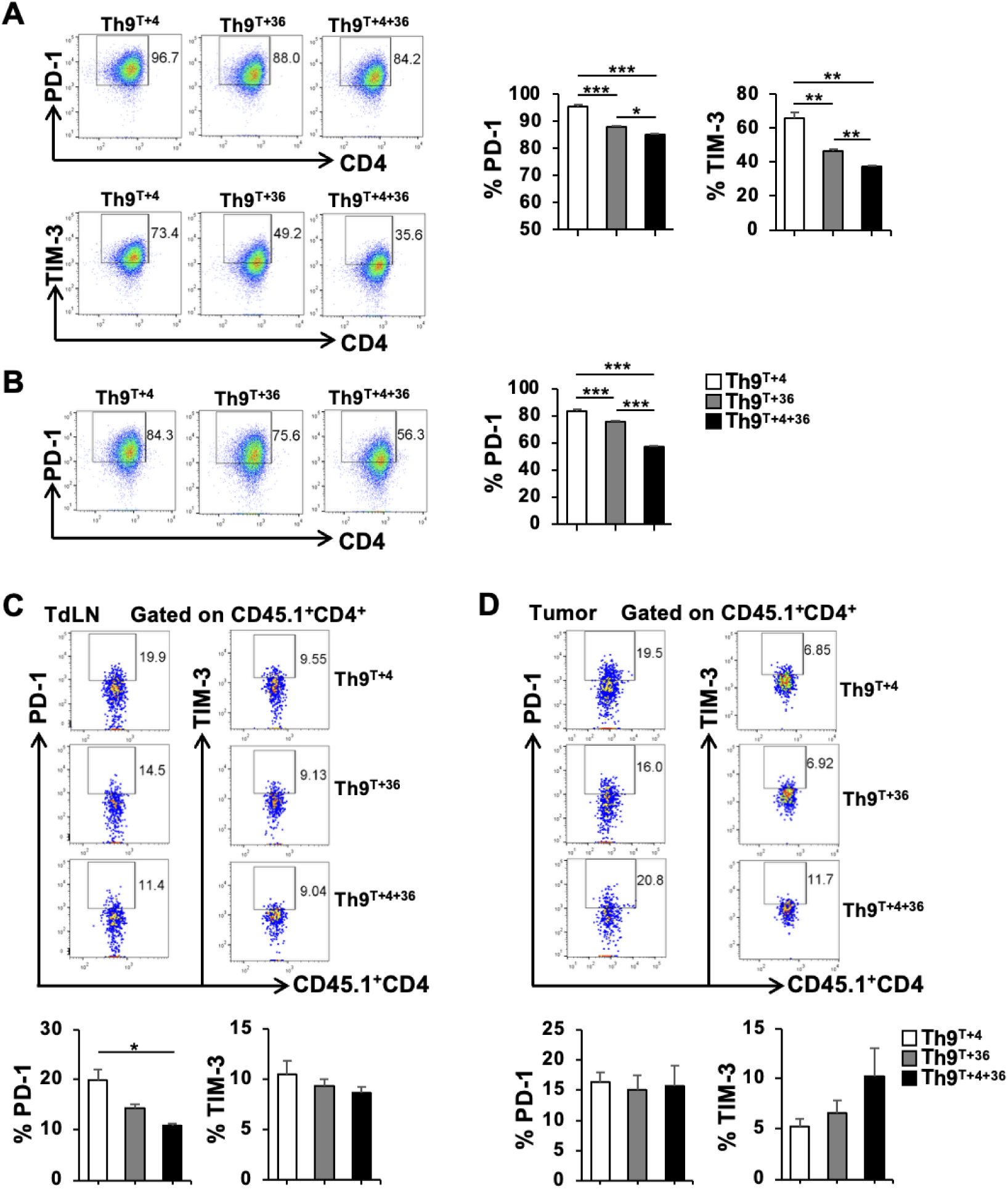
IL-36γ downregulates PD-1 and TIM-3 expression on *in vitro*-polarized Th9 subsets. **(A)** Naïve CD4^+^ T cells were activated with plate-bound anti-CD3/CD28 mAbs (each 5 μg/ml) in the presence of IL-2 (20 ng/ml) and cultured under Th9^T+4^, Th9^T+4+36^ or Th9^T+4+36^ conditions for 2 days. Representative flow cytometry plots of PD-1 and TIM-3 (left). Quantitative analysis of PD-1 and TIM-3 frequencies (right). **(B)** OVA_323-339_-specific CD4^+^ T cells from OT-II mice were activated with cognate peptide-pulsed APCs and polarized under Th9^T+4^, Th9^T+4+36^ or Th9^T+4+36^ conditions. Representative flow cytometry plots of PD-1 (left). Quantitative analysis of PD-1 frequency (right). **(C and D)** PD-1 and TIM-3 expression profiles on CD45.1^+^CD4^+^ T cells in TdLNs **(C)** and tumors **(D)** 10 days post-adoptive transfer. Representative flow cytometry plots of PD-1 and TIM-3 (up). Quantitative analysis of PD-1 and TIM-3 frequencies (below). One-way ANOVA was employed for data analysis **(A-D)**. \**p* < 0.05, \*\**p* < 0.01, \*\*\**p* < 0.001.

**Figure S7.**
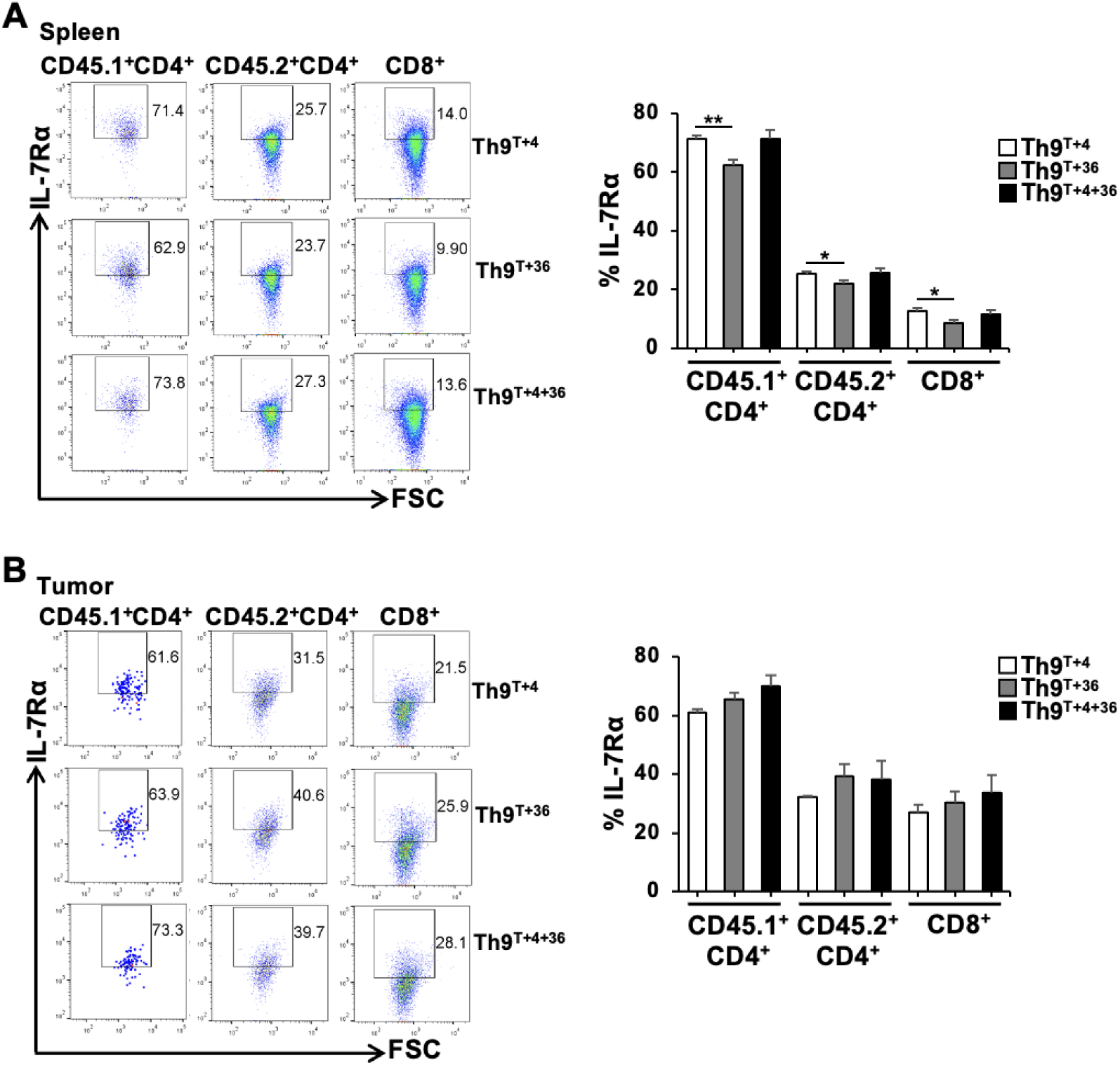
Quantification of IL-7Rα in adoptively transferred CD4^+^ T, host-derived CD4^+^ T and CD8^+^ T cells in spleens and tumors. **(A)** Flow cytometric analysis of IL-7Rα expression on in splenic T-cell subsets as indicated on day 20 post-ACT (left, representative flow cytometry plots; right, quantitative data). **(B)** Flow cytometric analysis of IL-7Rα expression in tumor-infiltrating T-cell subsets as indicated on day 20 post-ACT (left, representative flow cytometry plots; right, quantitative data). Intergroup differences across three Th9 therapeutic cohorts were analyzed by one-way ANOVA for indicated T cell subsets **(A and B)**. \**p* < 0.05, \**p* < 0.01.

---

**Supplemental Table 1.**
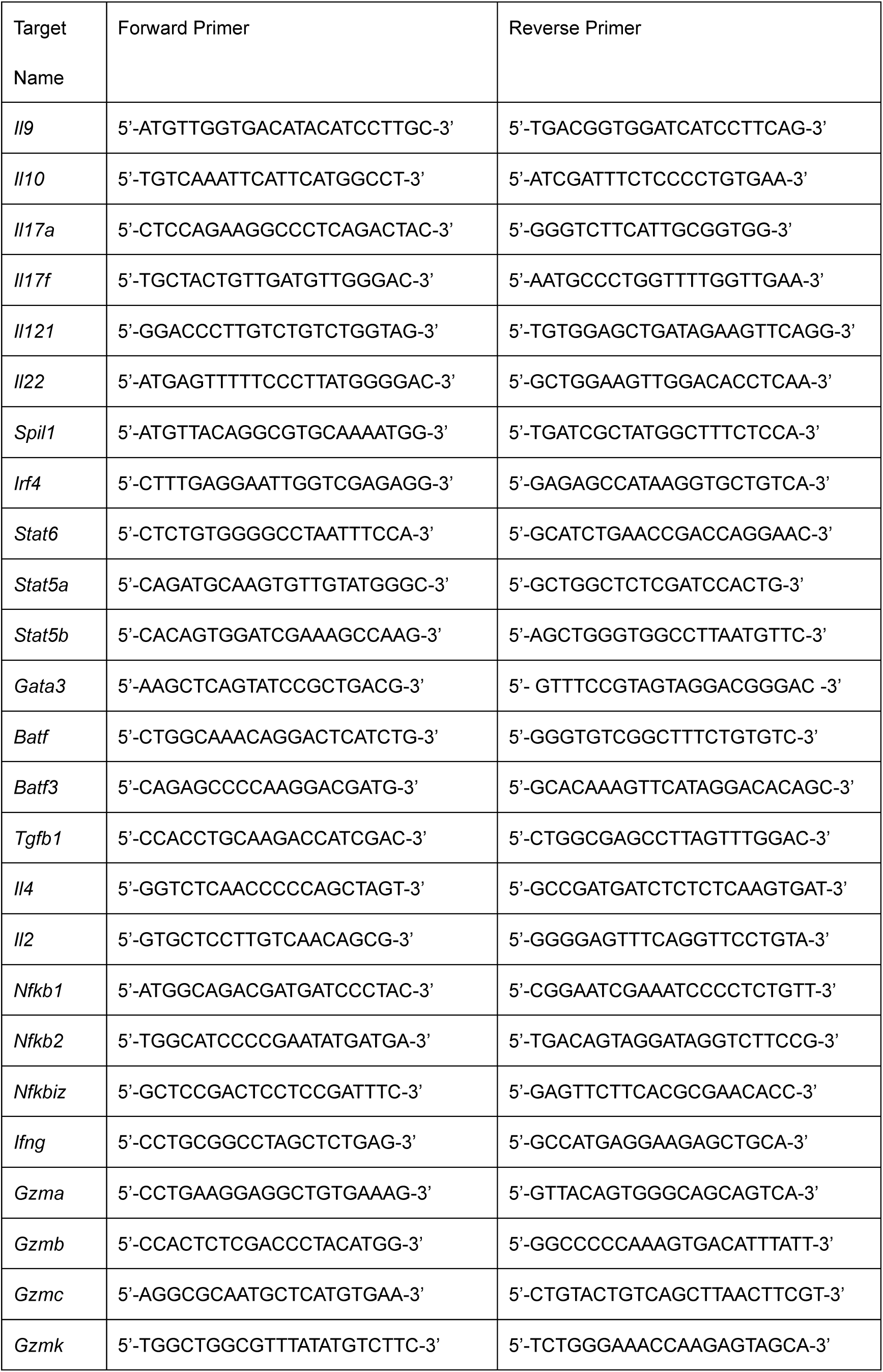
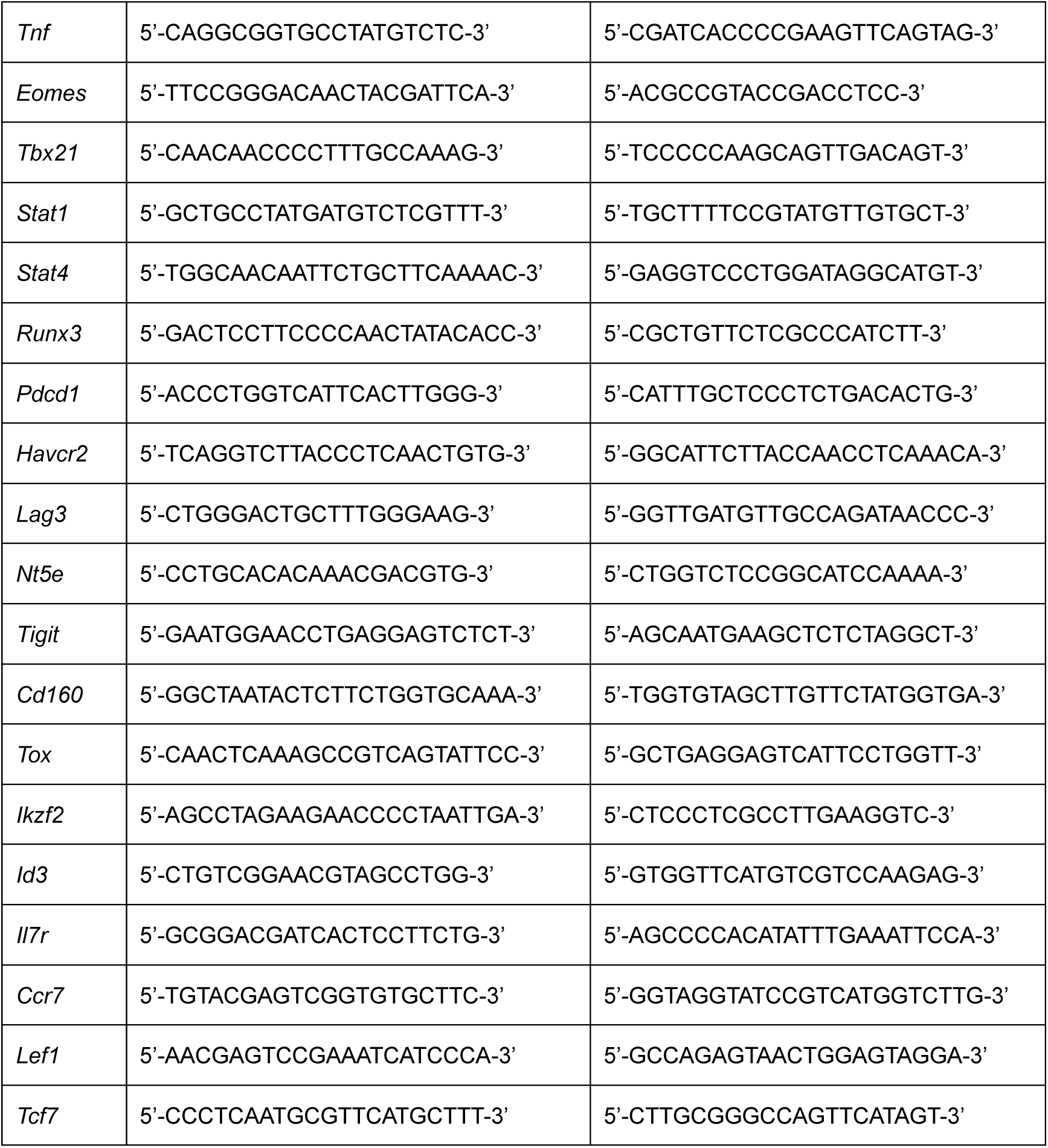
Primer sequence for qPCR.

**Supplemental Table 2.**
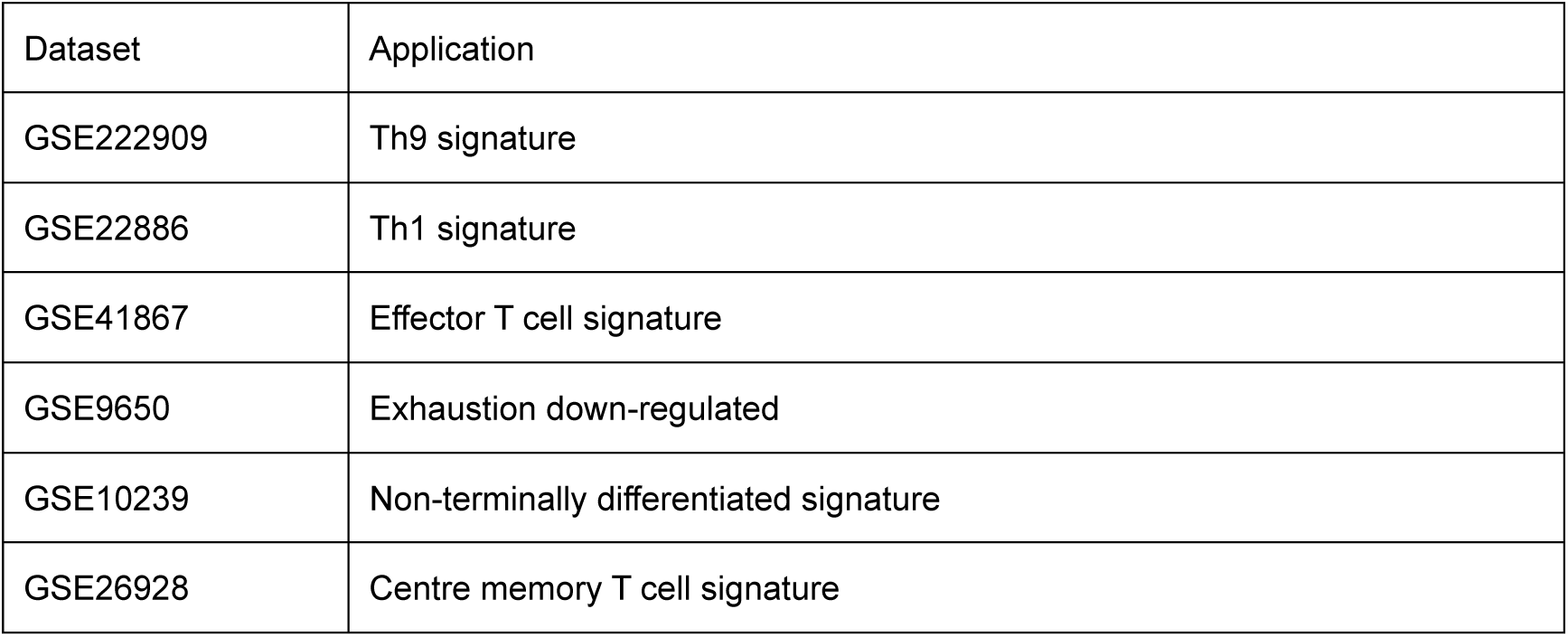
Public datasets analyzed.

